# A 3’UTR-derived small RNA modulates the life-cycle of the cholera toxin-encoding filamentous phage, CTXϕ

**DOI:** 10.64898/2025.12.02.691856

**Authors:** Anne Lippegaus, James R.J. Haycocks, Eoghan O’Driscoll, Marcel Sprenger, Kerstin Thriene, Elke-Martina Jung, Malte Siemers, Sebastian Krautwurst, David C. Grainger, Kai Papenfort

**Author notes:** To whom correspondence should be addressed: Kai Papenfort.

## Abstract

Bacteriophages (phages) are well known to be one of the major driving forces in bacterial evolution. This also applies to virulent microorganisms, such as the major human pathogen *Vibrio cholerae*, whose pathogenic potential and epidemic proliferation largely depends on the interaction with environmental phages. Specifically, integration of the CTXϕ phage genome into the first chromosome of *V. cholerae* also introduced the *ctxAB* genes, encoding the primary toxin responsible for the severe acute diarrheal disease, cholera. Whereas the mechanisms underlying CTXϕ-associated horizontal gene transfer and transcriptional control of the *ctxAB* genes have been intensively studied over the past years, post-transcriptional regulation affecting the CTXϕ life-cycle has not been documented. Here, we report the discovery and characterization of the CisR small RNA (sRNA) that is produced from the 3’UTR (untranslated region) of the *prtV* gene and inhibits the expression of the CTXϕ-encoded *cep* mRNA. CisR-mediated repression of *cep* involves Hfq-assisted base-pairing of the two transcripts and results in reduced CTXϕ production under stress conditions. We further demonstrate that transcription of *prtV-cisR* requires both the master quorum-sensing regulator HapR and CRP, a global regulator of carbon metabolism. Taken together, our work provides evidence that *V. cholerae* employs sRNA-mediated post-transcriptional gene regulation to coordinate CTXϕ activation with both cell density and nutrient availability.

**SIGNIFICANCE STATEMENT:** The integration of the CTXϕ phage genome, which carries the *ctxAB* toxin genes, is essential for cholera pathogenesis in humans. While transcriptional control of CTXϕ and *ctxAB* has been well-studied, post-transcriptional mechanisms remain unexplored. Here, we identify and characterize CisR, a small RNA derived from the 3′ untranslated region of *prtV*, which inhibits the CTXϕ-encoded *cep* mRNA through Hfq-dependent base-pairing. CisR-mediated regulation limits phage production under stress conditions and is co-regulated by the quorum-sensing factor HapR and the metabolic regulator CRP. Our findings reveal a new RNA-based mechanism linking CTXϕ phage activation to cell density and nutrient status of *V. cholerae*.

## INTRODUCTION

Cholera remains a significant health challenge in numerous regions of the developing world, accounting for an estimated ∼2.8 million cases and ∼90,000 fatalities annually (1). The causative agent, *Vibrio cholerae*, is ubiquitously present in marine environments, however, many *V. cholerae* strains are non-pathogenic or only cause sporadic instances of gastroenteritis (2). Pathogenicity of *V. cholerae* hinges on the acquisition of various virulence factors, particularly cholera toxin (CT) and the toxin-coregulated pilus (TCP). CT induces profuse watery diarrhea, a major contributor to cholera-related mortality and its rapid spread, while TCP facilitates colonization of the small intestine (3). The genes encoding CT, *ctxAB*, are carried in the genome of a lysogenic filamentous phage CTXϕ and genomic analyses of epidemic *V. cholerae* strains suggest multiple independent events of toxigenic conversion in the history of cholera (2, 4, 5).

Since its discovery (6), the CTXϕ life-cycle has been investigated in great detail. After entering *V. cholerae* cells, the single-stranded genome of CTXϕ can either integrate into the chromosome of *V. cholerae* or convert into double-stranded DNA to facilitate rolling-circle replication and transcription of genes crucial for phage replication and morphogenesis (7). The lysogenic phase of CTXϕ is mainly regulated by the phage repressor protein, RstR, which inhibits the key phage mobilization genes, *rstA* and *rstB*. Activity of RstR is supported by the host-encoded LexA protein (the master regulator of SOS response) and thereby links CTXϕ activation to DNA damage (8). In addition, RstR activity is counteracted by RstC, an antirepressor protein located on the RS1 satellite phage (9). The core components of CTXϕ consist of seven genes: *ctxAB* (encoding CT subunits A and B) and five structural genes (*cep*, *psh*, *g^IIICTX^*, *ace*, *zot*), which play crucial roles in phage morphogenesis and assembly (10). Similar to many other filamentous phages, CTXϕ secretion does not involve cell lysis, but rather depends on the type II secretion system (T2SS) of the host cell (11).

Besides LexA, several other host-encoded proteins have been reported to interact with CTXϕ at the genomic or regulatory level (12, 13). In this context, regulation of *ctxAB* regulation has been a major focus, reflecting the pivotal role of CT in cholera pathogenesis (14). Expression control of the *ctxAB* operon involves the ToxT transcriptional regulator that is encoded on another horizontally-acquired genomic element, called VPI-1 (*Vibrio* pathogenicity island 1). ToxT activity is modulated by bile and fatty acids (15, 16) and, besides *ctxAB*, controls the transcription of additional virulence-related genes, including the TarA and TarB sRNAs. Whereas TarA has been reported to regulate glucose uptake (17), TarB inhibits the expression of the *tcpF* virulence factor and has been documented to participate in phage defense functions (18–20).

Both, TarA and TarB, belong to the large group of Hfq-binding sRNAs that have been studied in various pathogenic bacteria, including *V. cholerae* (21). Hfq functions as an RNA chaperone that promotes sRNA stability and facilitates base-pairing of sRNAs with target mRNAs (22, 23). A single sRNA usually controls multiple target genes and in some cases sRNAs have been documented to rival transcription factors with respect to the number of regulated targets (24). Thereby, sRNAs can promote stress responses and virulence, coordinate metabolic fluxes, and support overall cellular homeostasis (25–28). Traditionally, sRNA genes have been discovered in intergenic regions of the genome, however, global transcriptome analyses have now revealed that sRNAs can also correspond to coding sequences, as well as the 5’ and 3’UTRs of mRNAs (29).

In this manuscript, we discovered a 3’UTR-derived sRNA from *V. cholerae* that is transcribed together with the *prtV* mRNA, encoding an extracellular protease involved in host-microbe interaction and biofilm formation (30, 31). Global RNA interactome studies revealed that the sRNA base-pairs with and inhibits the translation of multiple targets, including the *cep* mRNA of phage CTXϕ. Accordingly, we named the sRNA, CisR (<u>C</u>TXϕ inhibiting <u>s</u>mall <u>R</u>NA). Expression of the *prtV-cisR* transcript requires transcriptional activation by HapR and CRP, linking CisR accumulation to quorum sensing and general carbon metabolism, respectively. Mutation of *cisR* increased Cep protein levels under stress conditions and promoted CTXϕ production, suggesting modulation of CTXϕ phage levels. Taken together, our data indicate that the core-genome encoded CisR sRNA controls the production of the horizontally-acquired virulence gene that has a crucial role in the life-cycle of the CTXϕ phage.

## RESULTS

### Identification of differentially expressed sRNAs under virulence inducing conditions

Bacterial sRNAs are well known to be differentially expressed under various stress conditions, as well as when virulence gene expression is induced (32, 33). In *V. cholerae*, the TarA and TarB sRNAs have been reported to be activated by the ToxT transcriptional regulator, yet no systemic searches for virulence-associated sRNAs have been performed. To close this gap, we compared the gene expression profiles of wild-type *V. cholerae* cells cultivated in rich media (when virulence gene expression is repressed) with AKI growth conditions, a protocol known to activate virulence gene expression outside the human host (34). We confirmed efficient activation of virulence genes in AKI samples by quantifying the levels of the pathogenicity-associated *toxT* and *ctxA* mRNAs using qRT-PCR (quantitative real-time PCR). As expected, the expression of both transcripts was strongly upregulated under virulence-inducing conditions (Fig. S1). We next performed high-throughput-sequencing to determine the global gene expression changes associated with growth under AKI growth conditions in *V. cholerae*. These analyses revealed dozens of differentially expressed sRNAs and mRNAs, including known virulence-associated transcripts, such as *toxT* and the TarB sRNA (Fig. 1A). Other sRNAs displaying differential expression involved VssRNA24 (35), which was upregulated under AKI conditions, as well as the uncharacterized Vcr229 sRNA (36) displaying reduced expression (Figs. 1A-B). In addition, another uncharacterized sRNA, CisR (initially identified as Vcr096 (37)), was also upregulated under virulence-inducing conditions (Fig. 1A-B). The *cisR* gene is located at the 3’ end of *prtV* (Fig. 1C), encoding a Zn^2+^-binding extracellular protease, which is required by *V. cholerae* to infect *Caenorhabditis elegans* (30).

**Figure 1:**
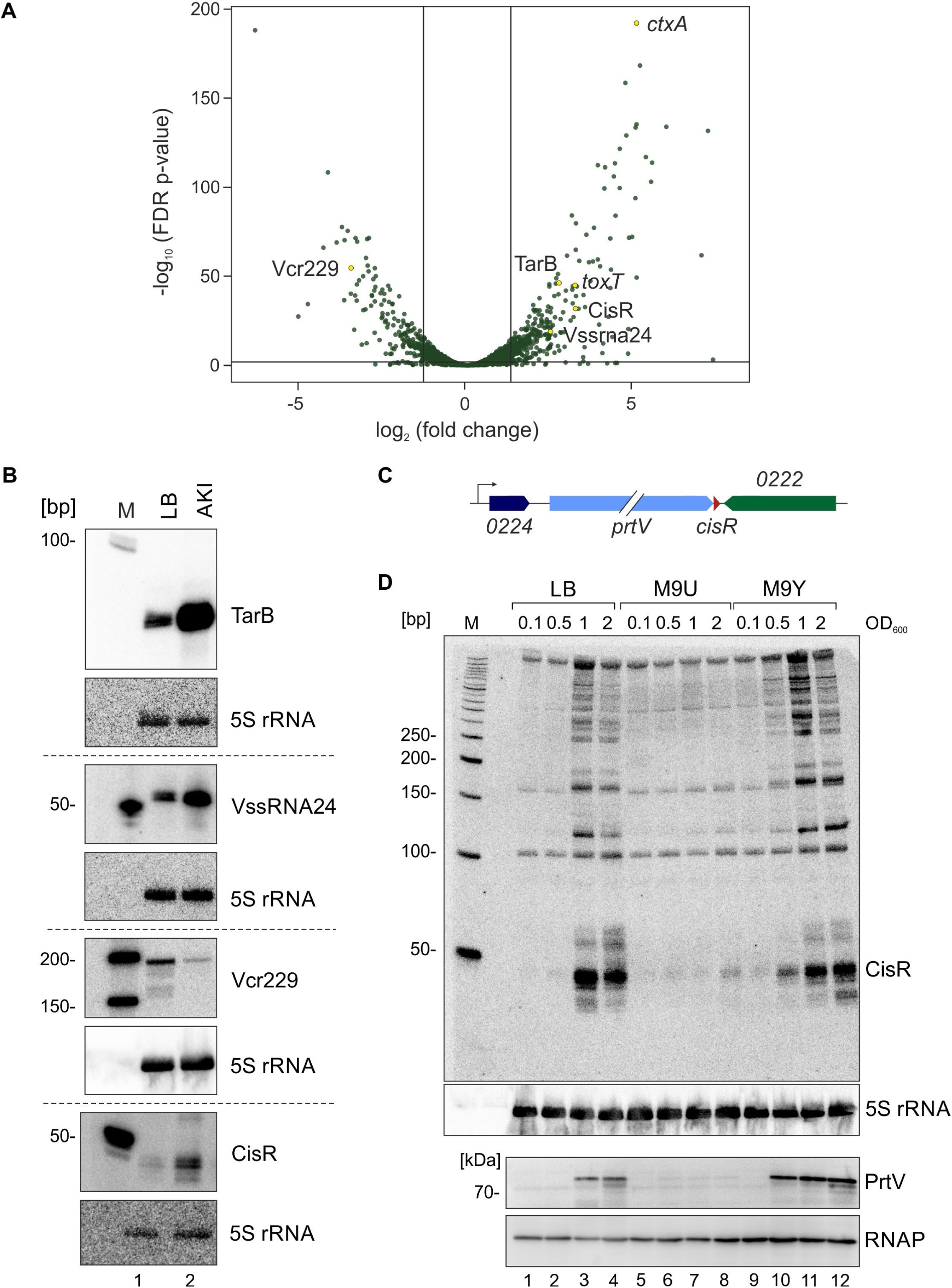
Transcriptomics in virulence inducing conditions. **(A)** Volcano plot of genome-wide transcript changes in response to virulence inducing conditions (AKI medium). Lines indicate cut-offs of differentially regulated genes at 3-fold and FDR-adjusted p-value≤0.01. Genes validated by qRT-PCR and Northern blotting are marked in yellow. **(B)** Validation of differentially regulated sRNAs by Northern blotting. RNA samples were obtained from *V. cholerae* wild-type cells cultivated in LB or AKI medium. sRNA levels were monitored by Northern blotting and probed with specific oligonucleotides for the indicated sRNAs. 5S ribosomal RNA served as a loading control. **(C)** Schematic representation of the genomic context of *cisR*. *cisR* is marked in red and *prtV* in light blue. **(D)** Expression of CisR and PrtV. *V. cholerae* wild-type cells were cultivated in LB, M9U (glucose) and M9Y (glycerol) medium. RNA and protein samples were collected at various stages of growth. Northern blot analysis was performed to determine CisR levels and Western Blot analysis was carried out to monitor PrtV protein levels. 5S ribosomal RNA and RNAP served as loading controls for Western and Northern blots, respectively.

To further investigate regulation of CisR in *V. cholerae*, we next probed its expression by Northern blotting. Here, we detected several *cisR*-specific bands and that the sRNA accumulates as a ∼50 nucleotide-long transcript (Fig.1D, top). In complex media (LB), expression of CisR was increased at high cell densities, whereas cultivation of *V. cholerae* in minimal media containing glucose as the sole carbon source resulted in strongly reduced CisR levels. In contrast, minimal media supplemented with glycerol activated CisR expression at high cell densities, suggesting that *cisR* synthesis is affected by carbon utilization and cell density. We also probed PrtV protein levels under same conditions using Western blot analysis. These data showed that PrtV and CisR display highly similar expression patterns (Fig.1D, bottom), suggesting that their expression might be coupled.

### CisR is produced from the 3’end of the *prtV* mRNA

Previous work on *V. cholerae* sRNAs indicated that a large fraction of these regulators are processed from longer transcripts (38, 39). To address this question for *prtV*-*cisR*, we first aligned the *cisR* sequences from various *Vibrio* species, showing that the region downstream of the *prtV* stop codon as well as segments of the 3’ UTR (including a Rho-independent terminator element) are highly conserved (Fig. S2A). In contrast, these analyses did not reveal any *cisR*-associated promoter element, supporting the hypothesis that the sRNA might be processed from the *prtV* mRNA. This idea was further supported by a previous study, indicating an RNase E cleavage sites at the 5’ end of *cisR* (40) (Fig. S2B).

To further test if accumulation of CisR in *V. cholerae* results from RNase E-mediated cleavage of the *prtV-cisR* transcript, we tested CisR expression in the absence of active RNase E. Although RNase E is essential in *V. cholerae* and many other bacteria (21), its function can be studied in a temperature-sensitive mutant strain (*rne*TS), which allows for the selective inactivation of RNaseE at non-permissive temperatures (40). Specifically, we collected total RNA samples from wild-type and the *rne*TS strain at permissive (30°C) and non-permissive (44°C) temperatures and compared CisR levels by Northern blotting. Our results showed that the mature CisR sRNA was not detected in the absence of RNase E (Fig. S2C), indicating that the endoribonuclease is indeed required for cleavage of the *prtV-cisR* transcript and CisR accumulation in the cell.

To further investigate the source of CisR expression in *V. cholerae*, we designed two plasmids: Plong, which includes a predicted promoter region upstream of the adjacent gene *vca0224* (encoding a hypothetical protein) (39), and a truncated version, Pshort, starting from the *prtV* start codon (testing for potential internal promoters in the *prtV* coding sequence) (Fig. S2D). We transferred both plasmids in *V. cholerae* Δ*cisR* cells and quantified CisR levels by Northern blotting. Comparison of the CisR-specific signals with *V. cholerae* wild-type cells carrying an empty vector control showed increased CisR expression from plasmid Plong, whereas the levels of CisR in Pshort were down-regulated. Taken together, our data suggest that *cisR* is co-transcribed with *prtV* using a shared promoter element located upstream of *vca0224*.

### CisR expression is activated by the HapR and CRP transcription factors

To identify factors controlling *vca0224*-*prtV*-*cisR* expression in *V. cholerae*, we next investigated potential regulatory DNA sequences located upstream of the *vca0224* gene. Alignment of the relevant sequences from various *Vibrio* strains revealed conserved -35 and - 10 promoter elements, as well as putative binding sites that would match the consensus sites of the HapR and CRP transcriptional regulators (41, 42) (Fig. S3A). To investigate the roles of the two transcription factors on CisR expession, we constructed *V. cholerae* single-gene deletion mutants of *hapR* and *crp*, as well as a double mutant lacking both genes and measured CisR levels at various growth conditions using Northern blot analysis (Fig. 2A). When compared to wild-type cells, the *hapR* mutant displayed reduced CisR levels at all stages of growth, with residual levels detected under early stationary phase growth conditions (OD600 of 1.0). CisR levels were even further reduced in the *crp* mutant and when both transcriptional regulators were absent. We observed a similar regulatory pattern using a GFP-based transcriptional reporter of the promoter upstream of *vca0224* (Fig. 2B) and using qRT-PCR, we confirmed that expression of the *prtV* mRNA is also regulated by HapR and CRP (Fig. S3B). In summary, these data indicated that both HapR and CRP positively regulate *prtV*-*cisR* expression by controlling the promoter upstream of *vca0224*.

**Figure 2:**
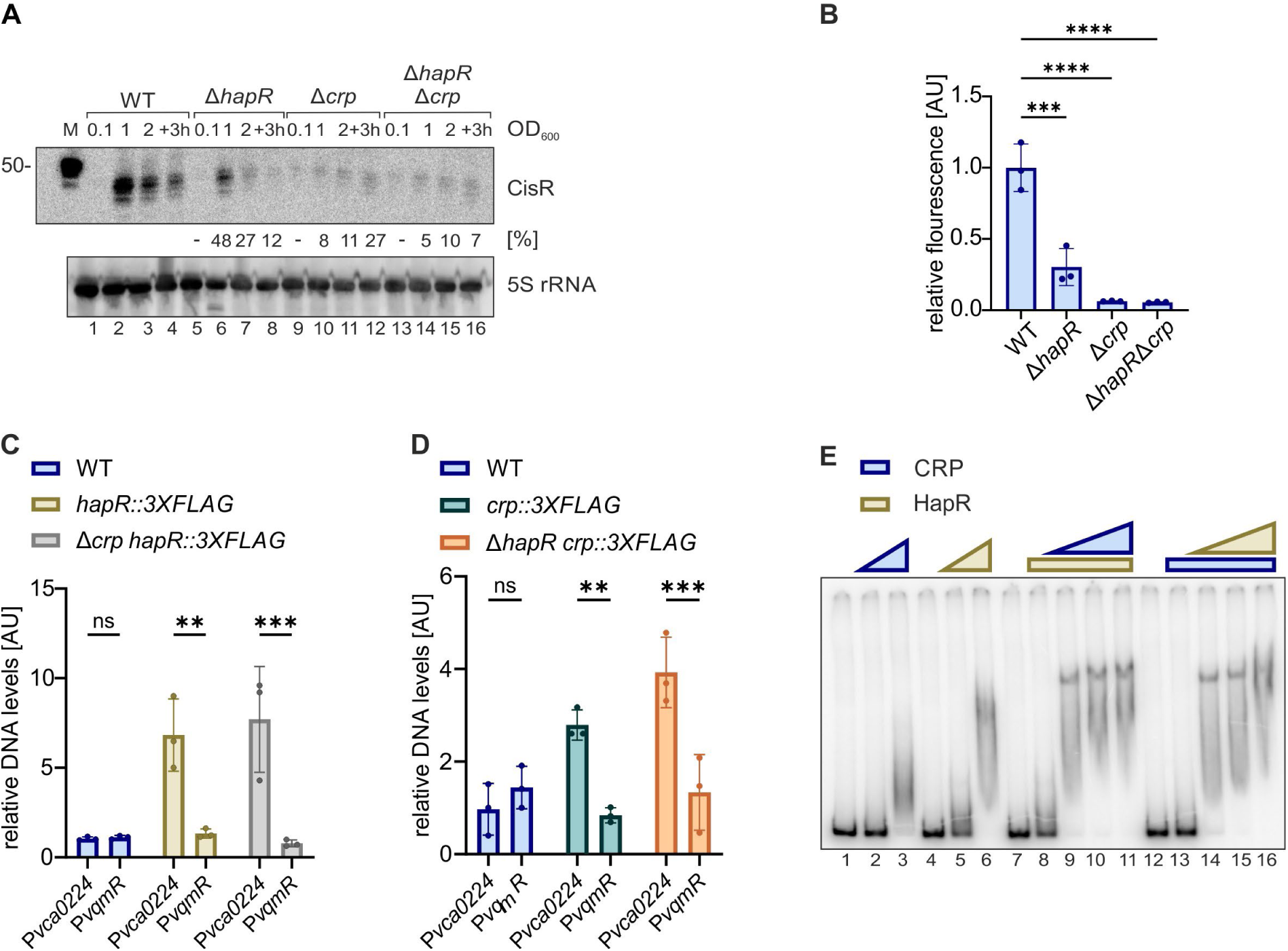
Transcriptional control of *cisR*. **(A)** Role of HapR and CRP for CisR levels. *V. cholerae* wild-type, Δ*hapR*, Δ*crp* and Δ*hapR* Δ*crp* were cultivated in LB medium and RNA samples were collected at different stages of growth. Northern blot analysis was performed to determine CisR levels. Probing for 5S ribosomal RNA served as loading control. **(B)** Regulation of the promoter located in front of *vca0224* (P*vca0224*). *V. cholerae* wild-type, Δ*hapR*, Δ*crp* and Δ*hapR* Δ*crp* carrying a GFP-based transcriptional reporter for *vca0224* (P*vca0224::GFP*) were cultivated in LB medium to OD600 of 1.0 and analyzed for fluorescence. *V. cholerae* wild-type were set to 1. Bars show mean of independent biological replicates ±SD, n=3. For statistical analyses, an ordinary one-way ANOVA with Dunnetts’s multiple comparison test was used (∗∗∗p ≤ 0.001, ∗∗∗∗p ≤ 0.0001). **(C)** ChIP analysis of HapR. *V. cholerae* wild-type, *hapR::3XFLAG* cells and *Δcrp hapR::3XFLAG* were cultivated to OD600 of 1.0 and subjected to chromatin immunoprecipitation (ChIP). Bar graphs show relative levels of *cisR* and *vqmR* promoters (P*vca0224* and P*vqmR*), determined by quantitative PCR and P*vca0224* levels in WT *V. cholerae* cells were set to 1. Data are presented as mean values of independent biological replicates ±SD, n=3. For statistical analyses, a two-way ANOVA with Šídák method for multiple comparisons was used (ns, not significant, ∗∗p ≤ 0.01, ∗∗∗p ≤ 0.001). **(D)** ChIP analysis of CRP. *V. cholerae* wild-type*, crp::3XFLAG* cells and Δ*hapR crp::3XFLAG* were cultivated to OD600 of 1.0 and subjected to chromatin immunoprecipitation (ChIP). Bar graphs show relative levels of the *cisR* and the *vqmR* promoters (P*vca0224* and P*vqmR*), determined by quantitative PCR and P*cisR* levels in WT *V. cholerae* were set to 1. Data are presented as mean values of independent biological replicates ±SD, n=3. For statistical analysis, a two-way ANOVA with Šídák method for multiple comparison was used (ns, not significant, ∗∗p ≤ 0.01, ∗∗∗p ≤ 0.001). **(E)** Electrophoretic mobility shift assay of HapR and CRP binding to P*vca0224*. Radiolabeled P*vca0224* transcript was incubated alone (lanes 1, 4, 7, 12), with increasing concentrations (0.5 µM to 2 µM) of purified HapR (lanes 2-3) and CRP (lanes 5-6) or with 0.5 µM HapR or CRP while increasing the other one from 0.5 µM to 2 µM (lanes 8-11 and 13-16). Complexes were separated on a native polyacrylamide gel and visualized by autoradiography.

To further investigate the individual roles of HapR and CRP for CisR expression, we generated *hapR* and *crp* complementation plasmids (p-*hapR* and p-*crp*, respectively) and tested their effect on CisR levels in *V. cholerae* wild-type cells and the relevant mutant strains by Northern blotting. Whereas both plasmids did not significantly change CisR expression in wild-type cells, p-*hapR* restored CisR expression in Δ*hapR* cells, but not in the Δ*crp* or Δ*crp*/hapR mutants (Fig. S3C). Similarly, p-*crp* restored CisR levels in the Δ*crp* mutant. Interestingly, Crp was also able to up-regulate CisR expression in the absence of *hapR*, suggesting that CRP has key regulatory function for *cisR* (and *prtV*) synthesis.

Following up on this hypothesis, we tested if binding of one transcription factor (*i.e.* CRP and HapR) to the *vca0224* promoter is interdependent or if both transcription factors can bind separately. To this end, we used chromatin immunoprecipitation (ChIP) assays. Specifically, we generated *V. cholerae* strains producing chromosomally tagged HapR::3XFLAG or CRP::3XFLAG proteins and performed immunoprecipitation using anti-FLAG antibodies. The co-purified DNA was analyzed by quantitative PCR. When compared to a non-tagged wild-type control, co-immunoprecipitation of HapR::3XFLAG enriched the *vca0224* promotor sequence ∼7-fold, while CRP::3XFLAG enriched the same region ∼3-fold (Fig. 2C-D), confirming that both HapR and CRP directly bind to the *vca0224* promoter *in vivo*. As a negative control, we tested co-immunoprecipitation of the unrelated *vqmR* promoter sequence, which showed no significant enrichment in either ChIP experiment. Importantly, binding of HapR to the *vca0224* promoter was observed in the absence of CRP, and likewise CRP binding occurred in the absence of HapR. These data indicated that HapR and CRP can both interact with the *vca0224* promoter, which we also tested *in vitro*. Here, we performed EMSA (electrophoretic mobility shift assay) using radiolabeled *vca0224* promoter DNA together with purified CRP and HapR proteins. In accordance with our ChIP data, both proteins were able to bind the promoter DNA individually, however, highly cooperative when CRP and HapR were applied in combination (Fig. 2E). Taken together, these results align with our *in vivo* observations (Figs. 2A and S3C), confirming that full *prtV-cisR* expression requires both HapR and CRP.

To study a potential interplay of CRP and HapR during *vca0224* promoter binding, we also tested a CRP variant carrying a E55A substitution (CRPE55A). This mutation disrupts a previously described interaction between CRP and HapR, which is required for binding of the *V. cholerae murQP* promoter (41). EMSA studies were conducted using both wild-type CRP and the CRPE55A variant in combination with HapR. Both, wild-type CRP and CRPE55A were able to bind the *vca0224* promoter region independently, and the presence of CRPE55A did not alter HapR’s ability to interact with the promoter (Fig. S3D). These findings suggest that the CRP E55 side chain is dispensable for binding at the vca0224 promoter and does not contribute to any direct interaction between CRP and HapR. In contrast, binding of either CRP or HapR might affect DNA topology and thereby facilitate binding of the other transcription factor. However, at this point, we cannot exclude the possibility that CRP and HapR could interact via a different side chain in CRP.

To better understand the simultaneous binding of CRP and HapR, we conducted *in vitro* DNase I footprinting assays to map their binding sites on the *vca0224* promoter. Specifically, radiolabeled DNA fragments containing the promoter fragment were incubated with purified CRP, HapR, or both, and then treated with DNase I. The resulting DNA fragments were analyzed on a denaturing gel to reveal areas of protection where the transcription factors prevented DNase I cleavage. Consistent with its typical mode of action, CRP binding produced a characteristic footprint with alternating regions of protection and hypersensitivity to DNase I digestion (41). These hypersensitive sites reflect CRP-induced DNA bending, which is a hallmark of CRP-DNA interactions (Fig. S3E). The CRP footprint spanned two distinct regions of the promoter: one between positions -247 and -225 bp, and another weaker footprint between positions -197 bp and -177 bp upstream of the *vca0224* transcriptional start site (Figs. S3A and S3E). HapR produced adjacent footprints (HapR I: -220 bp to -196 bp, HapR II: - 178bp to -152 bp) that did not overlap with the CRP-binding sites. In the presence of both CRP and HapR, the footprints persisted, suggesting that these factors do not compete for the same binding site. Instead, they occupy distinct yet adjacent sites on the promoter, supporting the above mentioned model proposing that binding of one transcription factor affects DNA topology, which might facilitate recruitment of the other transcription factor. Independent of the specific mode of action, our data show that transcription of *cisR* (and *prtV*) requires CRP and HapR and thus integrates information from carbon metabolism (CRP) and cell density (HapR).

### RIL-seq analysis reveals the Hfq-mediated RNA interactome under virulence conditions

We next aimed to study the regulatory role of CisR in *V. cholerae.* To this end, we employed RIL-seq (RNA-interaction-by-ligation-and-sequencing) (43, 44) to determine the global Hfq-mediated RNA-RNA interactome under virulence conditions. Using *V. cholerae* cells expressing Hfq::3XFLAG protein from its native chromosomal locus, we cultivated cells under AKI conditions (45). RIL-seq involves UV crosslinking and the ligation of RNA-RNA pairs bound by Hfq, followed by high-throughput sequencing of the resulting chimeric reads. This method thus provides a global map of Hfq-dependent interactions. A *V. cholerae* wild-type strain lacking the 3XFLAG epitope served as a negative control to ensure specificity of these interactions. Our analysis captured 1,395 RNA-RNA chimeras interacting with Hfq (Fig. 3A). Under virulence-inducing conditions, RIL-seq revealed that the vast majority (∼80%) of the CisR interactions involved RNA-RNA duplex with *cep* (core encoded pilus), which encodes a core structural element of the CTXϕ phage (46, 47). In contrast, under standard laboratory conditions (48), this interaction accounted only for ∼20% of CisR interactions (Fig. 3B). These findings highlight the dynamic nature of RNA networks in response to different growth conditions and suggested that CisR might regulate CTXϕ-encoded genes.

**Figure 3:**
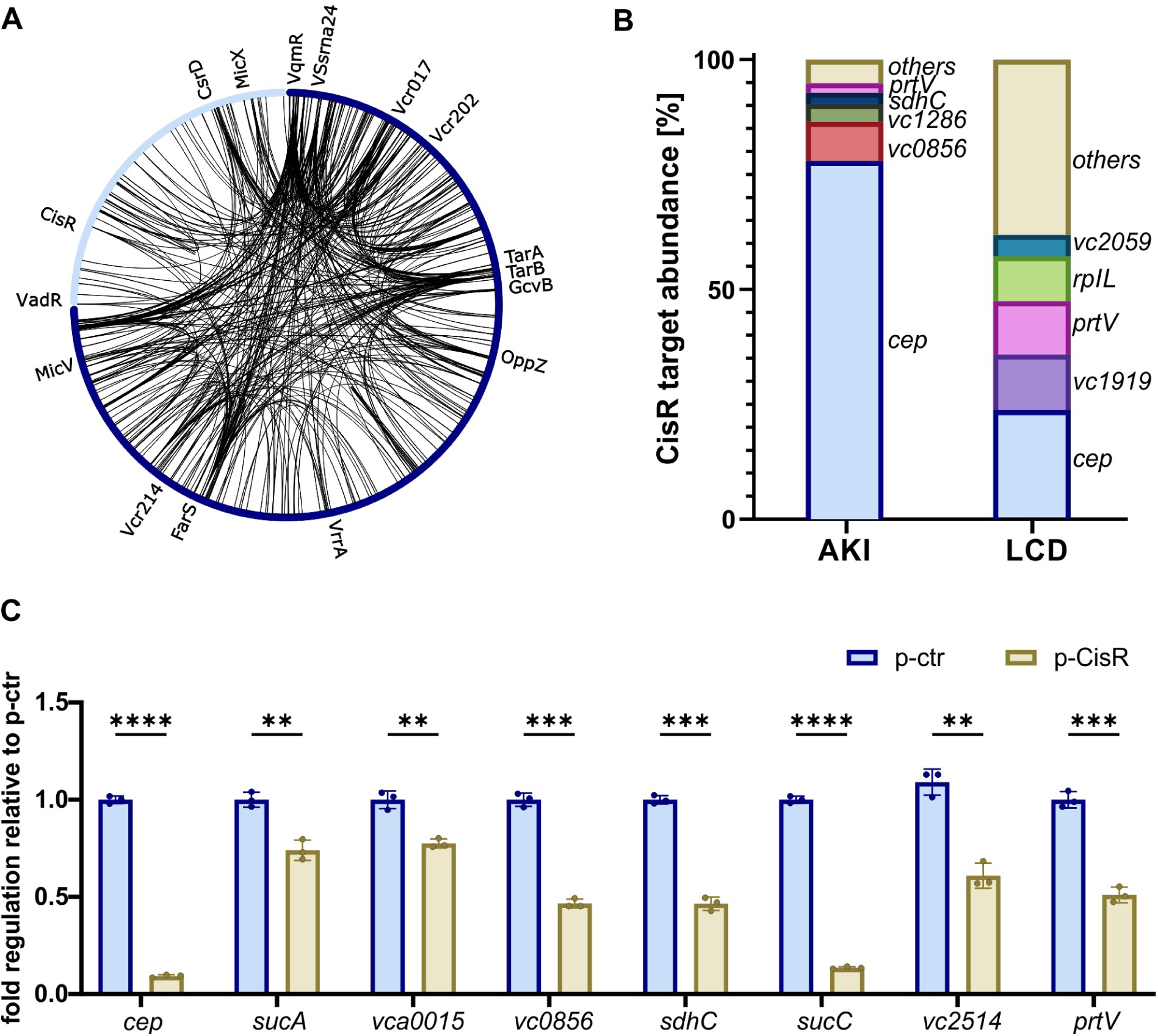
RIL-seq in *V. cholerae* under virulence inducing conditions. **(A)** Top 500 chimeras detected by RIL-seq under virulence inducing conditions. The first and the second chromosome are marked in dark and light blue, respectively. The circos plot was generated using the circos component of the Dash Bio package. **(B)** Comparison of CisR target abundance in two Hfq::3XFLAG RIL-seq datasets, virulence conditions (AKI) and low cell density (LCD) (48). The total amount of recorded CisR – mRNA interactions were set to 100% for each condition, respectively (LCD: 215 and AKI: 331). The top 5 interaction partners are named individually remaining interaction partners are summarized as others. **(C)** Validation of CisR targets predicted by RIL-seq. Translational GFP reporter fusions were co-transformed with a constitutive CisR expression plasmid (golden bars) or an empty control plasmid (blue bars) in *E. coli* Top10 cells. GFP production was measured and fluorophore levels from the control strains were set to 1. Error bars represent the SD of three independent biological replicates. Statistical significance was calculated using an unpaired t test (∗∗p ≤ 0.01, ∗∗∗p ≤ 0.001, ∗∗∗∗p ≤ 0.0001).

To confirm the regulatory role of CisR, we employed a GFP-based reporter system (49, 50) to monitor post-transcriptional control of eight potential targets (selected based on their abundance in RIL-seq experiments). In this system, one plasmid contains the 5’ untranslated region (5’ UTR) and the sequence encoding the first 20 amino acids of the target genes fused to *gfp*, whereas the sRNA is expressed from a second plasmid. In both cases, transcription is driven from constitutive promoters, thus specifically monitoring post-transcriptional gene regulation. We validated stable CisR expression in *E. coli* (Fig. S4A) and using the ChimericFragments workflow (51) together with the IntaRNA algorithm (52), we were able to predict RNA duplex formation of CisR with the putative targets (Figs. S4B-H). Indeed, this strategy allowed us to validate regulation of all eight targets by CisR, of which *cep* showed the strongest repression (Fig. 3C). Additional targets included *sucA* (2-oxoglutarate decarboxylase)*, vca0015* (hypothetical protein)*, vc0856* (DnaJ chaperone)*, sdhC* (succinate dehydrogenase subunit)*, sucC* (succinyl-CoA synthetase subunit)*, vc2514* (UDP-N-acetylglucosamine enolpyruvyl transferase) and also *prtV*, indicating that CisR could also act as a feedback regulator. Of note, a similar regulatory mechanism has previously been observed for the 3’UTR-derived OppZ and CarZ sRNAs in *V. cholerae* (53).

### CisR base-pairs with the *cep* mRNA encoded by CTXΦ

The above results suggested that the *cep* mRNA is the main target of CisR under virulence-inducing conditions and that its expression is post-transcriptionally repressed by the sRNA (Figs. 3B-C). To pinpoint the base-pairing position between CisR and *cep*, we probed the predicted RNA duplex by introducing point mutations at three critical positions (Fig. 4A). While mutation of CisR only mildly affected its expression (Fig. S5A), it strongly reduced the ability of the sRNA to inhibit the *cep::gfp* reporter (Fig. 4B). Likewise, mutation of *cep* abrogated repression by CisR, however, combination of both mutated CisR and *cep* variants restored regulation. These results confirmed the predicted RNA duplex and given that CisR interacts with the *cep* ribosome binding site, we predict that base-pairing results in inhibition of translation initiation at the *cep* mRNA.

**Figure 4:**
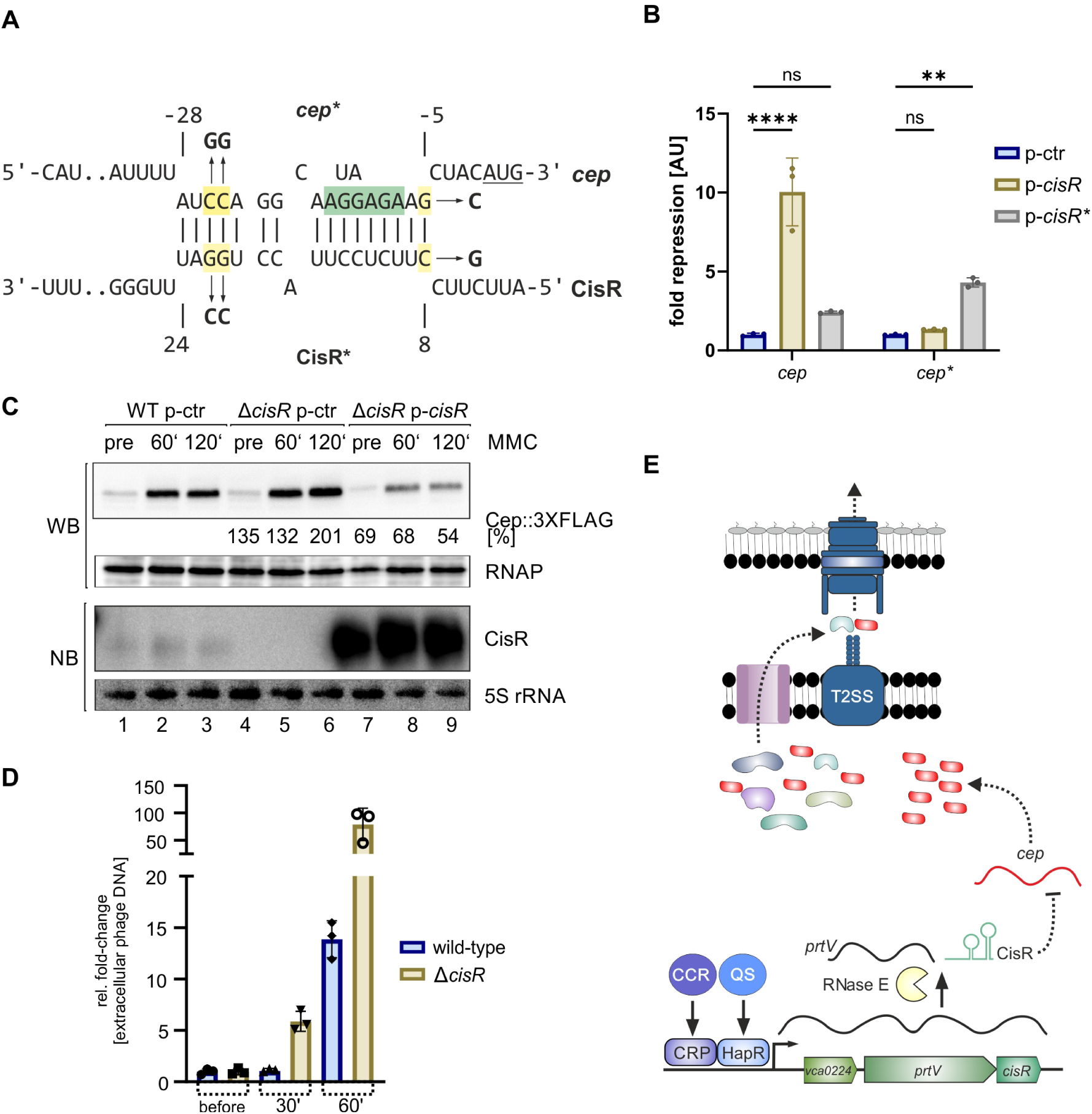
CisR inhibits CTXφ colonization factor. **(A)** Predicted base-pairing regions between CisR and *cep* by ChimericFragments (81). SD-sequence, start codon and point mutations used in (B) are highlighted. **(B)** Compensatory mutation CisR and *cep*. Translational GFP reporter fusion of *cep*/ *cep\** were co-transformed with constitutive CisR/ CisR* expression plasmids or an empty control plasmid in *E. coli* Top10 cells. GFP production was measured, fluorophore levels from the control strains were set to 1 and the fold repression was plotted. For statistical analyses, a two-way ANOVA with Dunnett’s multiple comparison test was used (ns, not significant, ∗∗p ≤ 0.01, ∗∗∗∗p ≤ 0.0001). **(C)** Effect of MMC on CisR and Cep. Protein and RNA samples of *V. cholerae cep::3XFLAG* p-ctr, Δ*cisR cep::3XFLAG* p-ctr and Δ*cisR cep::3XFLAG* p-*cisR* were collected in AKI medium pre, 60 min and 120 min post treatment with MMC (250 ng/mL). Protein levels of Cep::3XFLAG were monitored by Western blotting and CisR RNA levels were monitored by Northern blotting. RNAP and 5S ribosomal RNA served as loading controls for Western and Northern blot, respectively. **(D)** RstC-mediated activation of CTXϕ. Extracellular CTXϕ DNA levels were measured in cell-free supernatants of *V. cholerae* wild-type and Δ*cisR* cells carrying a pBAD-RstC plasmid using qPCR. DNA levels were quantified before and 30 and 60 minutes after RstC activation and normalized to the chromosomal *hfq* gene to control for potential cell lysis. The levels of CTXϕ DNA before pBAD induction were set to 1 and the relative production of extracellular CTXϕ phage DNA in wild-type and Δ*cisR* cells was plotted. **(D)** Regulatory model for CisR-mediated regulation of *cep*. Expression of the *vca0224-prtV-cisR* transcript is activated by the HapR and CRP transcription factors by quorum sensing (QS) and carbon catabolite repression (CCR). The transcript is processed by RNase E to release the CisR sRNA, which base-pairs with the *cep* mRNA to inhibit its translation. We speculate that high Cep levels might interfere with the type II-mediated secretion of other proteins in *V. cholerae*.

The Cep protein constitutes the main major coat protein of CTXϕ and is required for virion morphogenesis (6, 54). We therefore asked if CisR-mediated repression of *cep* also affects the production of CTXΦ when phage replication is induced. To this end, we cultivated *V. cholerae* under AKI conditions and used mitomycin C (MMC) to prompt LexA-mediated CTXΦ activation (8, 55). Using qRT-PCR, we quantified *cep* mRNA levels in wild-type, Δ*cisR*, and CisR over-expression strains. In the absence of MMC, we observed only basal *cep* expression and *cisR* mutation or over-expression only modestly affected *cep* levels (Fig. S5B). In contrast, MMC treatment strongly activated *cep* expression (>60-fold) and this effect was further enhanced in cells lacking *cisR* (>100-fold). In accordance with our hypothesis, over-expression of CisR reduced *cep* expression when compared to wild-type and Δ*cisR* cells. To corroborate these findings at the protein levels, we added a 3XFLAG epitope tag to the chromosomal *cep* gene and monitored Cep::3XFLAG production in the presence and absence of MMC and/or CisR. Analogous to *cep* mRNA levels, Cep::3XFLAG levels were drastically increased by MMC treatment and this effect was more pronounced in the Δ*cisR* mutant, when compared to isogenic wild-type cells (Fig. 4C). As expected, over-expression of CisR inhibited Cep::3XFLAG expression in the absence and presence of MMC. Taken together, our data indicate that CisR plays a regulatory role in modulating Cep levels during phage induction and thus might affect the CTXϕ life-cycle.

To test the latter, we harnessed the activity of the antirepressor RstC, which inhibits the phage repressor RstR and activates the transcription of genes required for CTXϕ production (9). Specifically, we cloned the *rstC* gene under the control of the inducible pBAD promoter and quantified CTXϕ phage DNA in cell supernatants using quantitative PCR (qPCR). As expected RstC activation resulted in increased CTXϕ phage DNA levels, however, this effect was strongly pronounced in Δ*cisR* cells when compared to the isogenic wild-type strain (Fig. 4D). Taken together, our data suggest that CisR limits CTXϕ production by inhibiting Cep synthesis.

## DISCUSSION

Filamentous phages play a crucial role in the evolution and pathogenicity of *Vibrio* species, particularly *Vibrio cholerae* (54, 56). The CTXϕ phage, which carries the cholera toxin (*ctxAB*) and zonula occludens toxin (*zot*), transforms non-toxigenic into toxigenic strains, facilitating pathogen dissemination through diarrhea (13, 57). Given the importance of the CTXϕ phage for the evolution and life-cycle of *V. cholerae*, the mechanisms underlying phage genome integration and replication have been studied in great detail over the past few years (10, 47). In contrast, less is known about the interplay of the CTXϕ prophage with its *V. cholerae* host, focusing mainly on transcriptional regulation of the *ctxAB* toxin genes (58). In this study, we provide evidence that post-transcriptional regulation is also relevant for CTXϕ biology and its dissemination under stress conditions.

The Cep protein is a homolog of protein pVIII gene of filamentous coliphage Ff and related phages and constitutes the major coat protein of the CTXϕ (59, 60). Cep and pVIII play a crucial role in forming the viral capsid and it has been estimated that phage Ff requires ∼2,700 copies of pVIII protein per virion (61). Thus, reducing Cep levels in the cell, *e.g.* by CisR-mediated repression, might provide an efficient mechanism to inhibit CTXϕ genome packaging and limit phage propagation. This process could help to coordinate CTXϕ replication with the nutrient status of the cell and reduce the negative impact of CTXϕ activation under stress conditions.

Unlike tailed phages, the larger size of filamentous phages prevents their assembly inside the cell (59). Instead, the virion assembles at the bacterial cell envelope, where the maturing phage is actively released through the cell envelope without lysing the host cell. Specifically, CTXϕ employs the EspD secretin of the Type II secretion system (T2SS) for phage release (11). It is therefore possible that CTXϕ activation and the associated surge of Cep proteins in the cell blocks the T2SS-mediated export of other proteins causing membrane stress and reduced fitness (Fig. 4E). Of note, CTXϕ also uses T2SS to release CT into the host environment and the PrtV protease, which is co-produced with CisR (Figs. 1C-D), is exported through this pathway as well (62, 63). CisR also inhibits *prtV* expression (Fig. 3C), suggesting an additional level of feed-back regulation that could balance PrtV over-production in the context of stress or as a result of transcriptional bursting (53).

Transcription of *prtV-cisR* requires both HapR and CRP and also involves an additional hypothetical gene, *vca0224*, which is located upstream of *prtV* (Figs. 1C and S2D). It is currently not known if *vca0224* constitutes a translated open reading frame, however, homologous proteins have been annotated in various other *Vibrio* species (64). The relevant promoter upstream of *vca0224* contains binding sites for CRP and HapR (Fig. S2A) and binding of one of the transcription factors seems to facilitate the recruitment of the other (Figs. 2E and S3D). Co-regulation of transcription by CRP and HapR has previously been observed in *V. cholerae* and has been studied at the molecular level for the *murPQ* promoter (41). In contrast to the *vca0224* promoter (Fig. S3A), HapR and CRP bind overlapping DNA sequences at this promoter, which results in HapR-mediated inhibition of CRP-dependent transcription. In addition, binding of CRP and HapR at the *murPQ* promoter requires direct protein-protein interaction via the negatively charged E55 surface residue of CRP, which does not seem relevant for binding at the *vca0224* promoter (Fig. S3D). Although at this point we cannot exclude direct interaction of CRP and HapR at the *vca0224* promoter through alternative modes of binding, we speculate that cooperativity of CRP and HapR at this promoter might be facilitated by changes in DNA structure and topology. In summary, our data support that joint transcriptional regulation by CRP and HapR is common in *V. cholerae* (and likely other Vibrios, (65)), however, depending on how the promoter elements are organized, the interaction between the two transcription factors can either activate or repress gene expression.

Joint regulation of *cisR* by CRP and HapR could also provide important insights into potential additional physiological roles of the sRNA. Whereas HapR levels in *V. cholerae* are tightly regulated by quorum sensing (66), activation of CRP occurs in the absence of the preferred carbon source (*i.e.* glucose) through carbon catabolite repression (CCR) (67). Indeed, expression of CisR is strongly repressed at low cell densities or in the presence of glucose (Fig. 1D). This expression pattern might also provide a regulatory link to other CisR targets (*e.g. sucA*, *sdhC*, *sucC*, *vc2514*, and *vca0015*; Fig. 3C), as these have been previously associated with quorum sensing and/or carbon metabolism (68, 69). Interestingly, *cep* is the only horizontally-acquired gene in the list of CisR targets, which raises interesting questions about the establishment and evolution of the associated base-pairing interactions (70). Given that *cisR* transcription is linked to *prtV* (Fig. S2B) and that CisR also inhibits *prtV* (Fig. 3C), we speculate autoregulation of *prtV* might constitute a conserved CisR function, whereas regulation of *cep* was added to the regulon at a later evolutionary stage when the CTXϕ phage integrated into the *V. cholerae* genome. This hypothesis is supported by our conservation studies, showing that *cisR* is conserved in various *Vibrio* species (Fig. S2A), however, these species typically do not encode the CTXϕ phage. Of note, other interactions between horizontally-acquired genes and core genome-encoded sRNAs have been previously described (71). One example is the highly conserved SgrS sRNA from *Salmonella enterica*, which base-pairs with and inhibits the translation of the *sopD* virulence factor mRNA (72). Likewise, horizontally-acquired sRNAs, *e.g.* encoded on phages and prophages, have been shown to regulate host mRNAs (73). For instance, the VpdS and Lpr1 sRNAs from vibriophage VP882 and coliphage λ, respectively, both base-pair with host-encoded targets and thereby facilitate phage replication (20, 74). Taken together, these data indicate that sRNA-mediated control of phage life-cycles might be more prevalent than previously expected.

## METHODS

### Bacterial strains and growth conditions

Bacterial strains used in this study are listed in Supplementary Table S1. *V. cholerae* and *E. coli* cells were grown in LB, AKI, M9U and M9Y media, as specified. The wild-type strain *V. cholerae* C6706 was used throughout the study. Unless otherwise stated, standard laboratory conditions consisted of cultures grown in LB medium at 37 °C with an aeration at 200 rpm. For specific experiments, *V. cholerae* C6706 cells were grown in AKI medium with 1 hour of aeration (34) to induce CT expression *in vitro*. Cultures were compared to cells grown under standard laboratory conditions and harvested at a similar OD600 of 0.7. Chromosomal mutations in *V. cholerae* were generated using the pKAS32 suicide plasmid for allelic exchange (75). Plasmids were introduced into *V. cholerae* by RK2/RP4-based conjugal transfer from *E. coli* S17λpir plasmid donor strains (76). Where appropriate, media were supplemented with antibiotics at the following concentrations: 100 µg/mL ampicillin; 20 µg/mL chloramphenicol; 50 µg/mL kanamycin; 50 U/mL polymyxin B; 5mg/mL streptomycin.

### Plasmids and DNA oligonucleotides

All plasmids and DNA oligonucleotides used in this study are listed in Supplementary Tables S2 and S3, respectively.

### RNA isolation, Northern blot analysis, and qRT-PCR

Total RNA was prepared and blotted as described previously (77). Nylon-membranes (Sigma #15356) were hybridized with [^32^P] labelled DNA oligonucleotides at 42°C or with riboprobes at 63°C. Riboprobes were generated using the MAXIscript T7 Transcription Kit (ThermoFisher, scientific, #AM1312). Signals were visualized using a Amersham Typhoon phosphoimager (GE Healthcare) and quantified with GelQuant software (BiochemLabSolutions). Oligonucleotides for Northern blot analysis are provided in Supplementary Table S3. For qRT-PCR, total RNA was DNA-digested using TURBO^TM^ DNAse (ThermoFisher scientific, #AM2238). qRT-PCR was performed using the Luna® Universal One-Step RT-qPCR Kit (New England Biolabs, #E3005). *hfq* was used as reference gene. Oligonucleotides used in all qRT-PCR analyses are also provided in Supplementary Table S3.

### Fluorescence measurements

Fluorescence assays to measure GFP expression were conducted as previously described (49). *E. coli* Top 10 cells were cultivated overnight in LB medium. To measure promoter activity, *V. cholerae* strains carrying *Pvca0224*::GFP transcriptional reporter were cultivated in LB medium and samples were collected at indicated time-points. For all fluorescence measurements, three independent replicates were used for each strain. Cells were washed and resuspended in PBS and relative fluorescence was determined using a Spark 10 M plate reader (Tecan). Control samples not expressing fluorescent proteins were used to subtract background fluorescence.

### Western blot analysis

Experiments were performed as previously described (78). Denatured protein samples were separated using SDS-PAGE and transferred to PVDF membranes for Western blot analysis. 3XFLAG-tagged fusions were detected using anti-FLAG antibody (Sigma, #F1804). RnaPα served as a loading control and was detected using anti-RnaPα antibody (BioLegend, #WP003). Signals were visualized using a Fusion FX EDGE imager (Vilber) and band intensities were quantified using BIO-1D software (Vilber).

### RstC-mediated CTXϕ activation

*V. cholerae* wild-type and Δ*cisR* cells carrying pBAD::RstC were grown to early stationary phase (OD600 of 1.0) and phage DNA was isolated from cell supernatants before and after the addition of L-arabinose (0.2% final conc.). qPCR was employed to quantify extracellular CTXϕ phage DNA (*cep* gene). Cell lysis was controlled by also tested the levels of the chromosomally-encoded *hfq* gene. The relevant oligonucleotides used for qPCR are listed in Table S3.

### Transcriptomic analysis using RNA-seq

The *V. cholerae* wild-type strain was cultivated in AKI and LB media, as described above. Total RNA was isolated and DNase digested (ThermoFisher scientific, #AM2238). RNA integrity was assessed using an Agilent 2100 Bioanalyzer. cDNA libraries were prepared with the NEBNext® Multiplex Small RNA Library Prep Set for Illumina® (New England Biolabs, #E7300) and cDNA library quality was confirmed on an Agilent 2100 Bioanalyzer. Pooled cDNA libraries were sequenced on a NextSeq1000 system with a 100-nt read length in single-read mode. Demultiplexed raw reads were trimmed for quality and adaptor sequences, then mapped to the *V. cholerae* reference genome (NCBI accession numbers NC_002505.1, NC_002506.1) using CLC Genomics Workbench (Qiagen) with standard parameter settings. Fold enrichment under virulence inducing conditions was compared to standard laboratory conditions using the CLC “Differential Expression for RNA-Seq” tool. Genes with a fold change ≥ 3 and an FDR-adjusted p-value ≤ 0.01 were defined as differentially expressed.

### ChIP and quantitative PCR

*V. cholerae* wild-type, *hapR::3xFLAG*, Δ*crp hapR::3xFLAG*, *crp::3xFLAG*, and Δ*hapR crp::3xFLAG* strains were cultivated to OD600 of 1.0 in LB medium. ChIP experiments to determine transcription factor binding were performed as described previously (48). Quantitative PCR (qPCR) was performed using GoTaq® qPCR Master Mix (Promega, #A6002) and the CFX96 Real-Time PCR System (Bio-Rad), with *recA* as the reference gene. The oligonucleotides used for qPCR are listed in Supplementary Table S3.

### EMSA and DNase I footprint

Promoter DNA fragments were excised from plasmid pSR, end-labeled with γ^32^-ATP, and treated with T4 PNK (New England Biolabs, #M0201). EMSAs and DNase I footprinting experiments were performed as described previously (79)

### RIL-seq experiment

*V. cholerae* wild-type and *hfq::3XFLAG* strains were cultivated in triplicates in AKI medium. Cells were harvested after 1h of aeration. The experiment was performed as previously described (44), with modifications in the rRNA depletion. rRNA was depleted using rRNA-specific biotinylated probes (80). Data analysis and visualization was performed according to our previously published computational pipeline (81).

## Supporting information

Combined SI

## DATA, MATERIALS, AND SOFTWARE AVAILABILITY

Raw reads and read count data of the RNA-Seq experiments were deposited into the Gene Expression Omnibus repository under accession number GSE311502. Raw reads and RNA-RNA-interaction data of the RIL-Seq experiments were deposited into the Gene Expression Omnibus repository under accession number GSE311501. Documentation of the ChimericFragments is available at: https://github.com/maltesie/ChimericFragments

## ACKNOWLEDGMENTS

We thank Andreas Starick, Kristin Gluth and Yvonne Greiser for their excellent technical support. α-PrtV antibody was a generous gift from Sun Nyunt Wai (Umeå University). This work was supported by the DFG (CRC1127-3 – Project-ID 239748522 and EXC 2051 – Project-ID 390713860) and the European Research Council (CoG-101088027).

## AUTHOR CONTRIBUTIONS

AL, JR, DG, EO and KP designed the experiments; AL, EO, and JRJH performed the experiments; AL, JRJH, EO, MSp, MSi, DG, SK, EMJ, and KP analyzed data; and AL and KP wrote the manuscript.

