## Supplementary material for "A 3’UTR-derived small RNA modulates the life-cycle of the cholera toxin-encoding filamentous phage, CTXϕ": Combined SI

**This supplement contains:**

Figures S1-5

Tables S1-3

Supplemental References

**TABLE OF CONTENTS**

**Figure S1** Expression of *ctxA* and *toxT* under virulence inducing conditions

**Figure S2** RNase E-mediated processing of *cisR*

**Figure S3** Transcriptional control of P*vca0224*

**Figure S4** Interaction partners of selected CisR targets discovered by RIL-seq

**Figure S5** Induction of CisR with MMC

**Table S1** Strains used in this study

**Table S2** Plasmids used in this study

**Table S3** Oligonucleotides used in this study

### Supplementary Figures

**Figure S1**

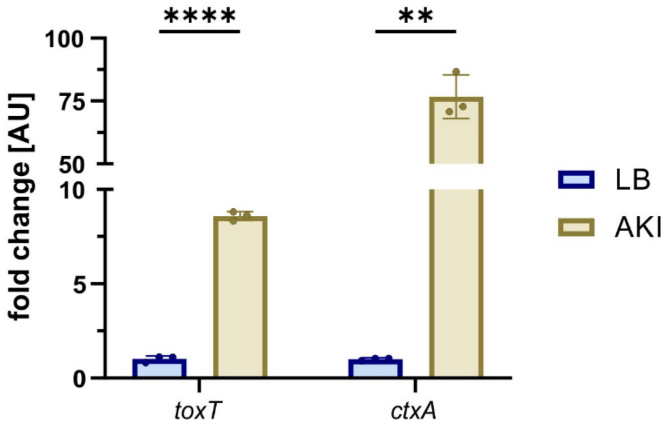


**Figure S1: Expression of *ctxA* and *toxT* under virulence inducing conditions**

*ctxA* and *toxT* mRNA levels are differentially expressed in virulence inducing conditions. RNA samples were obtained from *V. cholerae* wild-type cells were cultivated in LB and AKI medium. mRNA levels were determined by RT-qPCR. Data are presented as mean values of independent biological replicates ±SD, *n*=3. Statistical significance was calculated using an unpaired t test (**$p \leq 0.01$, ****$p \leq 0.0001$).

### Figure S2.

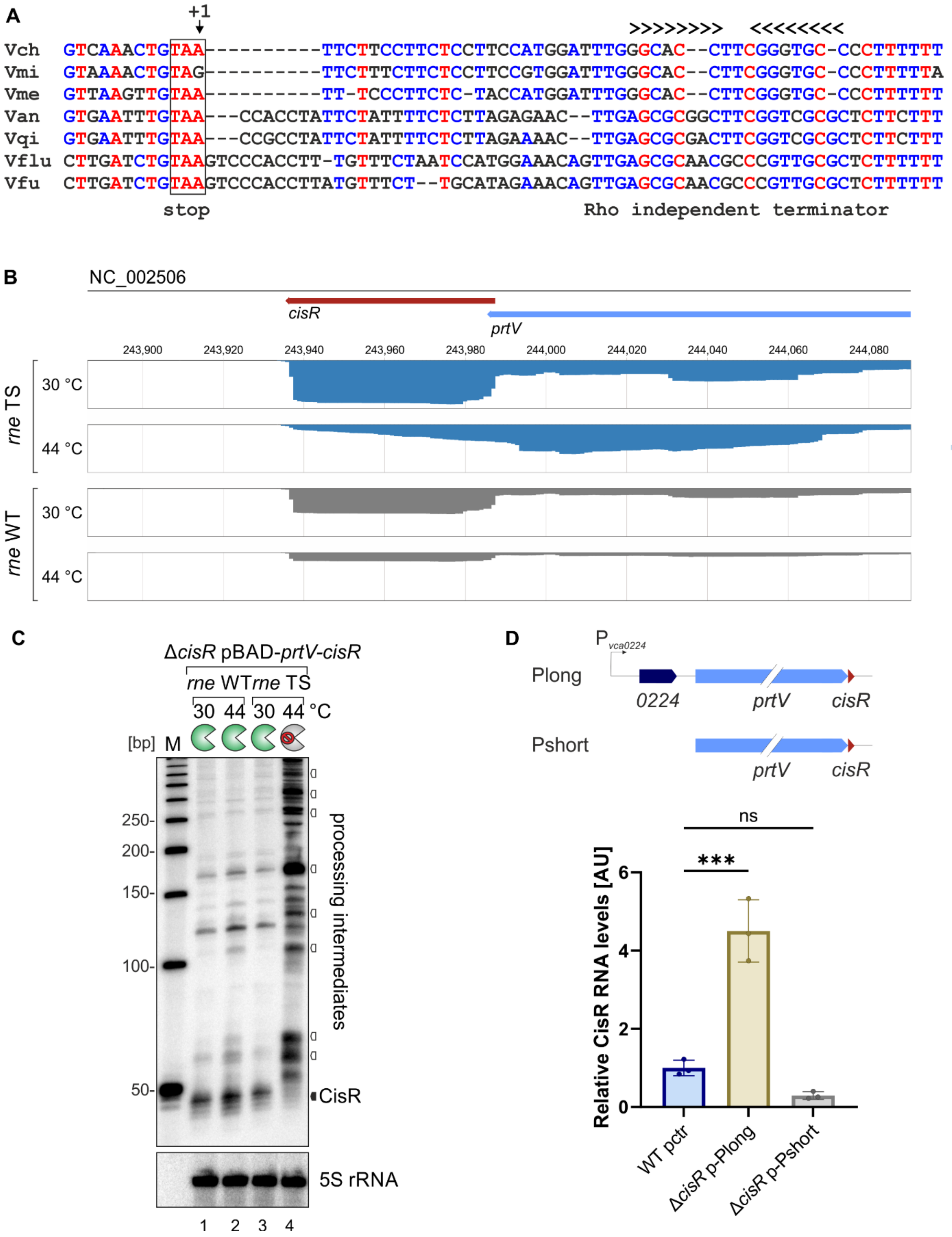

A
+1
>>>>>>>>   <<<<<<<<<
Vch GTCAAACTGTAA----------TTCTTCCTTCTCCTTCCATGGATTTGGGCAC--CTTCGGGTGC-CCCTTTTTT
Vmi GTAAAACTGTAG----------TTCTTTCTTCTCCTTCCGTGGATTTGGGCAC--CTTCGGGTGC-CCCTTTTTA
Vme GTTAAGTTGTAA----------TT-TCCCTTCTC-TACCATGGATTTGGGCAC--CTTCGGGTGC-CCCTTTTTT
Van GTGAATTTGTAA---CCACCTATTCTATTTTCTCTTAGAGAAC--TTGAGCGCGGCTTCGGTCGCGCTCTTCTTT
Vqi GTGAATTTGTAA---CCGCCTATTCTATTTTCTCTTAGAAAAC--TTGAGCGCGACTTCGGTCGCGCTCTTCTTT
Vflu CTTGATCTGTAAGTCCCACCTT-TGTTTCTAATCCATGGAAACAGTTGAGCGCAACGCCCGTTGCGCTCTTTTTT
Vfu CTTGATCTGTAAGTCCCACCTTATGTTTCT--TGCATAGAAACAGTTGAGCGCAACGCCCGTTGCGCTCTTTTTT
stop
Rho independent terminator
B
NC_002506
cisR
prtV
243,900
243,920
243,940
243,960
243,980
244,000
244,020
244,040
244,060
244,080
rne TS
30 °C
44 °C
rne WT
30 °C
44 °C
C
ΔcisR pBAD-prtV-cisR
rne WT
rne TS
30 44 30 44 °C
[bp]
M
250-
200-
150-
100-
50-
processing intermediates
CisR
5S rRNA
1 2 3 4
D
P vca0224
Plong
0224
prtV
cisR
Pshort
prtV
cisR
ns
***
Relative CisR RNA levels [AU]
0
2
4
6
WT pctr
ΔcisR p-Plong
ΔcisR p-Pshort

**Figure S2: RNase E-mediated processing of *cisR***

**(A)** Sequence alignment of *cisR* homologues in different *Vibrio* species. The *cisR* sequences were aligned using the Multalin tool (1). The stop codon of *prtV* is boxed and the Rho-independent terminator are indicated. *Vch, Vibrio cholerae; Vmi, Vibrio mimicus; Vme, Vibrio metoecus; Van, Vibrio anguillarum; Vqi, Vibrio qinghaiensis; Vflu, Vibrio fluvialis; Vfu, Vibrio furnissii.*

**(B)** Read-mappings of TIER-seq (2) to the *cisR* locus. The y-axes were set to the same scale. The positions in *V. cholerae* C6706 genome are indicated above the sequencing tracks. *prtV* and *cisR* are annotated. The reads for *rne* WT are presented in grey and for *rne* TS in blue. Temperatures are indicated on the left side of the tracks.

**(C)** Influence of RNase E on CisR levels. *V. cholerae* Δ*cisR* carrying either a wild-type (*rne* WT) or temperature-sensitive RNase E (*rne* TS) allele and the pBAD-*prtV*-*cisR* vector were cultivated at 30°C in LB medium. AT $OD_{600}$ of 1.0, cultures were split and kept at 30°C or shifted to 44°C for 30min. Expression of *prtV*-*cisR* was induced using L-arabinose (0.2%, 30min) and CisR levels were monitored by Northern blot. Probing for 5S ribosomal RNA served as loading control.

**(D)** Promotor identification of *cisR*. Quantification Northern Blot, Promotor identification. For statistical analysis, an ordinary one-way ANOVA with Dunnet's multiple comparison test was performed (ns, not significant, $***p \leq 0.001$).

**Figure S3.**

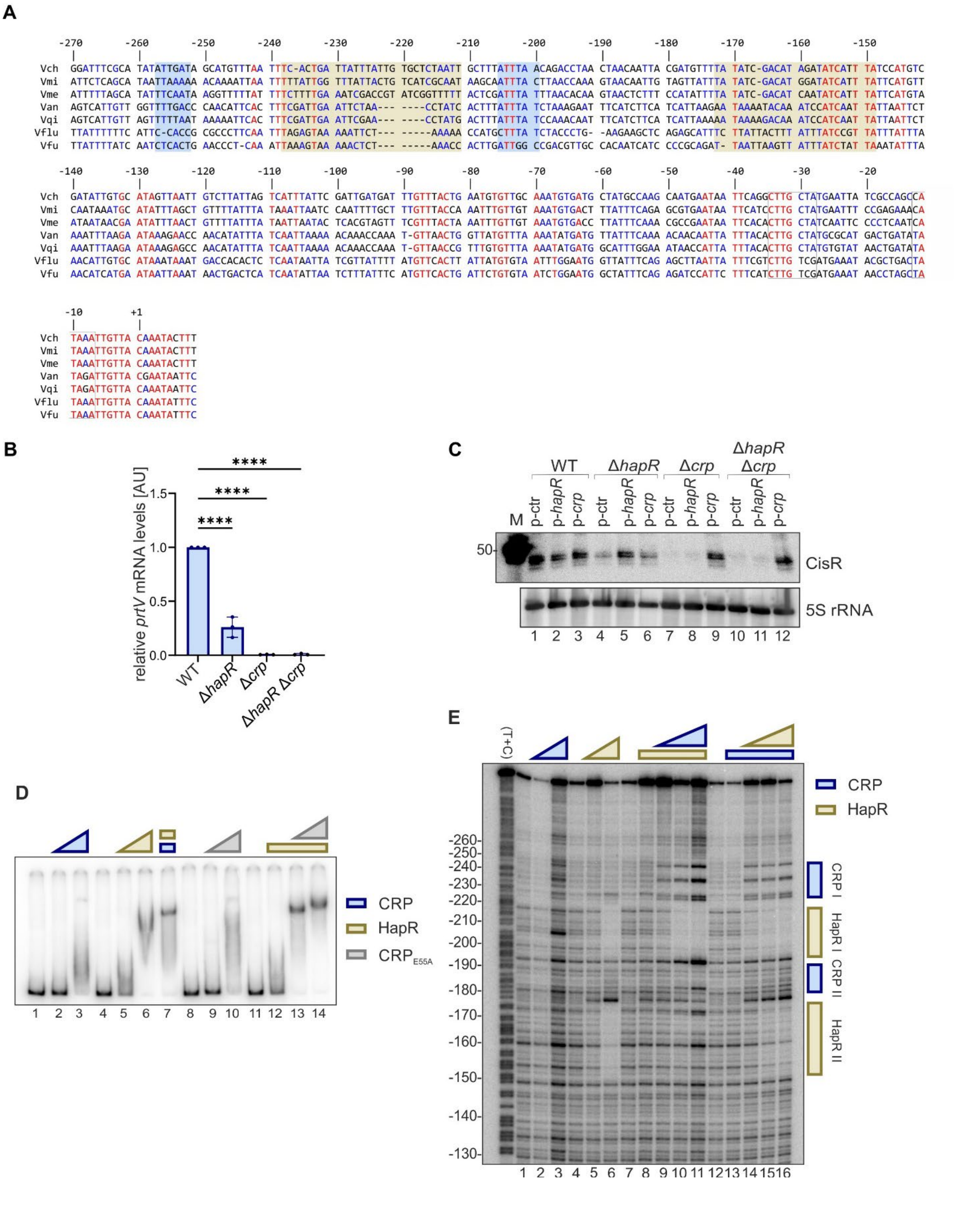
A
Vch GGATTTCGCA TATATTGATA GCATGTTTAA TTTC-ACTGA TTATTTATTG TGCTCTAATT GCTTTATTTA ACAGACCTAA CTAACAATTA CGATGTTTTA TATC-GACAT AGATATCATT TATCCATGTC
Vmi ATTCTCAGCA TAATTAAAAA ACAAAATTAA TTTTTATTGG TTTATTACTG TCATCGCAAT AAGCAATTTA CTTAACCAAA GTAACAATTG TAGTTATTTA TATC-GACAT GGATATCATT TATTCATGTT
Vme ATTTTTAGCA TATTTCAATA AGGTTTTTAT TTTCTTTTGA AATCGACCGT ATCGGTTTTT ACTCGATTTA TTTAACACAA GTAACTCTTT CCATATTTTA TATC-GACAT CAATATCATT TATTCATGTA
Van AGTCATTGTT GGTTTTGACC CAACATTCAC TTTCGATTGA ATTCTAA--- ----CCTATC ACTTTATTTA TCTAAAGAAT TTCATCTTCA TCATTAAGAA TAAAATACAA ATCCATCAAT TATTAATTCT
Vqi AGTCATTGTT AGTTTTTAAT AAAAATTCAC TTTCGATTGA ATTCGAA--- ----CCTATG ACTTTATTTA TCCAAACAAT TTCATCTTCA TCATTAAAAA TAAAAGACAA ATCCATCAAT TATTAATTCT
Vflu TTATTTTTTC ATTC-CACCG CGCCCTTCAA TTTAGAGTAA AAATTCT--- -----AAAAA CCATGCTTTA TCTACCCTG- -AAGAAGCTC AGAGCATTTC TTATTACTTT ATTTATCCGT TATTTATTTA
Vfu TTATTTTATC AATCTCACTG AACCCT-CAA ATTAAAGTAA AAACTCT--- -----AAACC ACTTGATTGG CCGACGTTGC CACAATCATC CCCGCAGAT- TAATTAAGTT ATTTATCTAT TAAATATTTA
Vch GATATTGTGC ATAGTTAATT GTCTTATTAG TCATTTATTC GATTGATGAT TTGTTTACTG AATGTGTTGC AAATGTGATG CTATGCCAAG CAATGAATAA TTCAGGCTTG CTATGAATTA TCGCCAGCCA
Vmi CAATAAATGC ATATTTAGCT GTTTTATTTA TAAATTAATC CAATTTTGCT TTGTTTACCA AATTTGTTGT AAATGTGACT TTATTTCAGA GCGTGAATAA TTCATCCTTG CTATGAATTT CCGAGAAACA
Vme ATAATAACGA ATATTTAACT GTTTTATTTA TAATTAATAC TCACGTAGTT TCGTTTACTA AATTTGTTGT AAATGTGACC TTATTTCAAA CGCCGAATAA TTCACACTTG CTATCAATTC CCCTCAATCA
Van AAATTTAAGA ATAAAGAACC AACATATTTA TCAATTAAAA ACAAACCAAA T-GTTAACTG GTTATGTTTA AAATATGATG TTATTTCAAA ACACAATTTA TTTACACTTG CTATGCGCAT GACTGATATA
Vqi AAATTTAAGA ATAAAGAGCC AACATATTTA TCAATTAAAA ACAAACCAAA T-GTTAACCG TTTGTGTTTA AAATATGATG GCATTTGGAA ATAACCATTA TTTACACTTG CTATGTGTAT AACTGATATA
Vflu AACATTGTGC ATAAATAAAT GACCACACTC TCAATAATTA TCGTTATTTT ATGTTCACTT ATTATGTGTA ATTTGGAATG GTTATTTCAG AGCTTAATTA TTTCGTCTTG TCGATGAAAT ACGCTGACTA
Vfu AACATCATGA ATAATTAAAT AACTGACTCA TCAATATTAA TCTTTATTTC ATGTTCACTG ATTCTGTGTA ATCTGGAATG GCTATTTCAG AGATCCATTC TTTCATCTTG TCGATGAAAT AACCTAGCTA
Vch TAAATTGTTA CAAATACTTT
Vmi TAAATTGTTA CAAATACTTT
Vme TAAATTGTTA CAAATACTTT
Van TAGATTGTTA CGAATAATTC
Vqi TAGATTGTTA CAAATAATTC
Vflu TAAATTGTTA CAAATATTTC
Vfu TAAATTGTTA CAAATATTTC
B
relative prtV mRNA levels [AU]
WT
ΔhapR
Δcrp
ΔhapR Δcrp
C
WT
ΔhapR
Δcrp
ΔhapR Δcrp
M
p-ctr
p-hapR
p-crp
CisR
5S rRNA
D
CRP
HapR
CRP E55A
E
(T+C)
CRP
HapR
CRP I
HapR I
CRP II
HapR II

**Figure S3: Transcriptional control of P*vca0224***

**(A)** Alignment of *cisR* promoter sequence from various *Vibrio* species. The -35 box, -10 box and TSS (+1) are indicated. Putative HapR and CRP binding sites are highlighted in gold and blue, respectively. *Vch, Vibrio cholerae; Vmi, Vibrio mimicus; Vme, Vibrio metoecus; Van, Vibrio anguillarum; Vqi, Vibrio qinghaiensis; Vflu, Vibrio fluvialis; Vfu, Vibrio furnissii.*
**(B)** Role of HapR and CRP for *prtV* levels. *V. cholerae* wild-type, Δ*hapR*, Δ*crp* and Δ*hapR* Δ*crp* cells were cultivated in LB medium and RNA samples were collected at $OD_{600}$ of 1.0. *prtV* mRNA levels were determined by qRT-PCR. For statistical analysis, an ordinary one-way ANOVA with Dunnetts's multiple comparisons test was used (∗∗∗∗$p \leq 0.0001$).
**(C)** Role of HapR and CRP for CisR levels. *V. cholerae* wild-type, Δ*hapR*, Δ*crp* and Δ*hapR* Δ*crp* cells harboring either an empty control vector (p-ctr), a *hapR* overexpression plasmid (p-*hapR*) or a *crp* overexpression plasmid (p-*crp*) were cultivated in LB medium and RNA samples were collected at $OD_{600}$ of 1.0. Northern blot analysis was performed to determine CisR levels and probing for 5S ribosomal RNA served as loading control.
**(D)** Electrophoretic mobility shift assay of $CRP_{E55A}$ binding to P*vca0224*. Radiolabeled P*vca0224* transcript was incubated alone (lanes 1, 4, 8 and 11), with increasing concentrations from 0.5 µM to 2 µM of purified CRP (lanes 2-3), HapR (lanes 5-6) and $CRP_{E55A}$ (lanes 9-10). In lane 7 the P*vca0224* transcript was incubated with 0.5 µM HapR and CRP respectively and in lanes 12-14 with constant concentrations of 0.5 µM HapR while increasing the $CRP_{E55A}$ concentration from 0 µM to 2 µM. Complexes were separated on a native polyacrylamide gel and visualized by autoradiography.
**(E)** Binding locations of HapR and CRP upstream of *vca0224* were determined by DNase I footprinting. The pattern of DNase I cleavage in the absence of any proteins is shown in lanes 1, 4, 7 and 12. Radiolabeled P*vca0224* transcript is incubated with increasing concentration of CRP (lanes 2-3) and HapR (lanes 5-6) respectively. In lanes 8-11 the transcript was incubated with constant concentrations of HapR while CRP concentrations were increased and vice versa in lanes 12-16. Changes in the cleavage pattern due to the presence of CRP and/ or HapR are indicated at the right side of the gel by blue and golden boxes, respectively.

**Figure S4.**

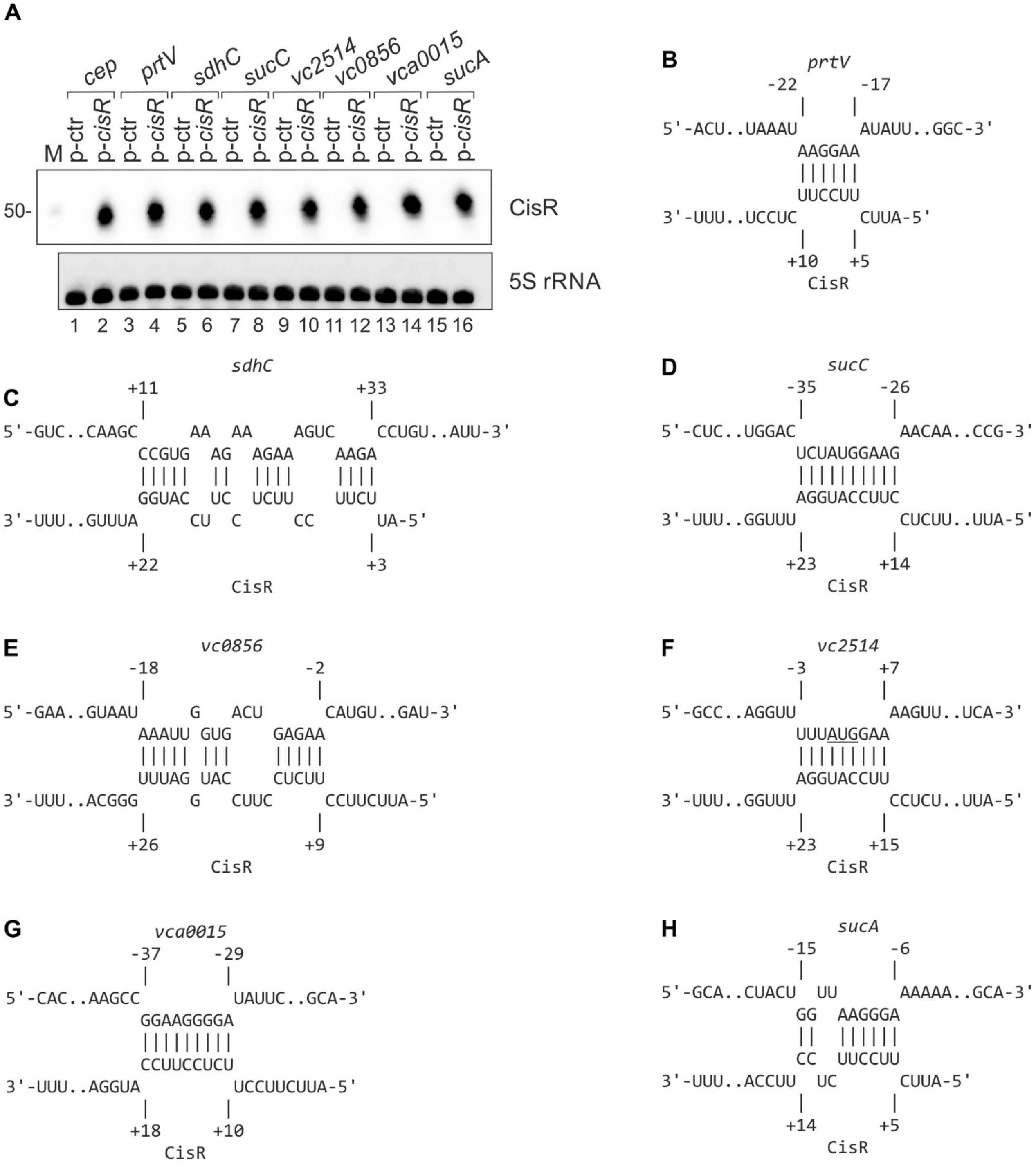


**Figure S4: Interaction partners of selected CisR targets discovered by RIL-seq**

**(A)** Plasmid-borne expression of *cisR* in *E. coli* Top10 cells. RNA samples are corresponding to Fig. 3C. CisR levels were determined by Northern blotting and probing for 5S ribosomal RNA served as loading control.

**(B)-(H)** Predicted base-pairing regions between CisR and its targets validated in Fig. 3C by ChimericFragments (3) and IntaRNA (4): *prtV* (B), *sdhC* (C), *sucC* (D), *vc0856* (E), *vc2514* (F), *vca0015* (G) and *sucA* (H). Start codons are underlined.

**Figure S5.**

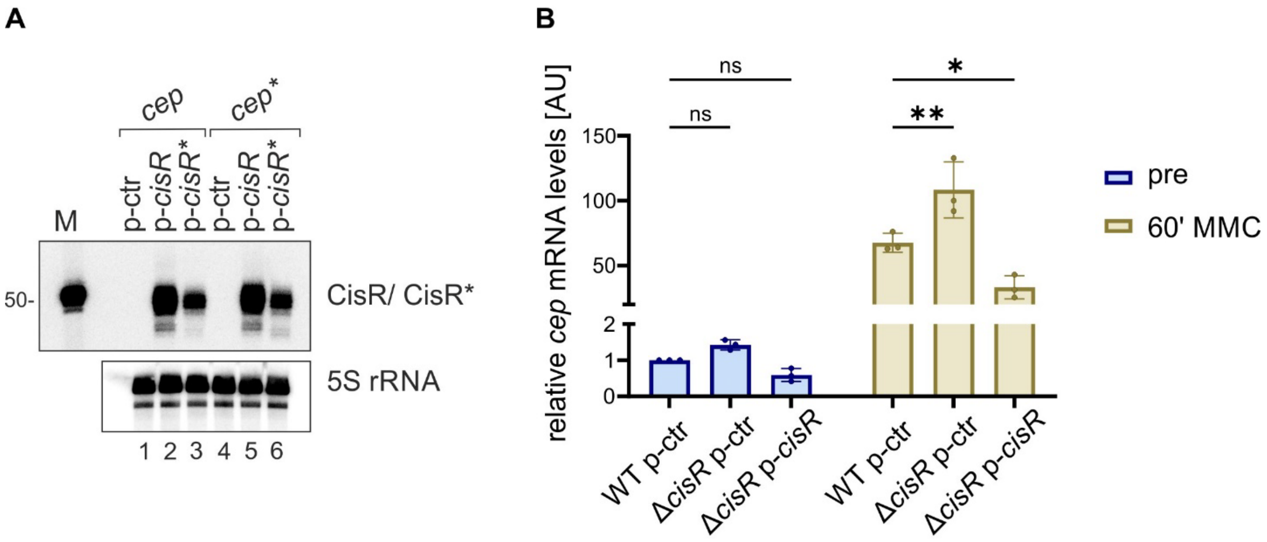


**Figure S5: Induction of CisR with MMC**

**(A)** Plasmid-borne expression of *cisR* and *cisR** in *E. coli* Top10 cells. RNA samples are corresponding to Fig. 4B. CisR/ CisR* levels were determined by Northern blotting and probing for 5S ribosomal RNA served as loading control.

**(B)** Effect of MMC on *cep* mRNA levels. RNA samples of *V. cholerae cep::3XFLAG* p-ctr, Δ*cisR cep::3XFLAG* p-ctr and Δ*cisR cep::3XFLAG* p-cisR were collected in AKI medium pre and 60 min post treatment with MMC (250 ng/mL). *cep* mRNA levels are determined by qRT-PCR. For statistical analysis, a two-way ANOVA with Tukey's multiple comparison test was used (ns, not significant, $*p \leq 0.05$, $**p \leq 0.01$).

### Table S1: Strains used in this study

| Strain | Relevant markers / Genotype | Reference / Source |
|---|---|---|
| ***V. cholerae*** | | |
| KPS-0014 | C6706 wild-type | (5) |
| KPS-0053 | C6706 Δ*hapR* | (6) |
| KPS-0995 | C6706 *hfq::3XFLAG* | (7) |
| KPVC-10966 | C6706 Δ*hapR* Δ*crp* | This study |
| KPVC-10984 | C6706 Δ*crp* | (8) |
| KPVC-12925 | C6706 *hapR::3XFLAG* | (9) |
| KPVC-13484 | C6706 *crp::3XFLAG* | This study |
| KPVC-14998 | C6706 Δ*cisR* | This study |
| KPVC-10141 | C6706 *rne*-3071 | (2) |
| KPVC-15015 | C6706 *rne*-3071 Δ*cisR* | This study |
| KPVC-15080 | C6706 *cep::3XFLAG* | This study |
| KPVC-15081 | C6706 Δ*cisR cep::3XFLAG* | This study |
| KPVC-15469 | C6706 Δ*hapR crp::3XFLAG* | This study |
| KPVC-15538 | C6706 Δ*crp hapR::3XFLAG* | This study |
| ***E. coli*** | | |
| Top10 | *F- mcrA Δ(mrr-hsdRMS-mcrBC) φ80lacZΔM15 ΔlacX74 nupG recA1 araD139 Δ(ara-leu)7697 galE15 galK16 rpsL(StrR) endA1 λ-* | Invitrogen |
| S17λpir | *ΔlacU169 (ΦlacZΔM15), recA1, endA1, hsdR17, thi-1, gyrA96, relA1, λpir* | New England Biolabs |

### Table S2: Plasmids used in this study

| Plasmid trivial name | Plasmid Stock Name | Relevant fragment | Comment | Origin, marker | Reference |
|---|---|---|---|---|---|
| pXG10-sfGFP | pXG10-sfGFP | lacZ'::sfGFP | Template plasmid for translational reporter | pSC101*, $Cm^R$ | (10) |
| pXG10-*vc1461* (*cep*) | pAL42 | 5'UTR + 20 aa of *vc1461* | Translational GFP reporter | pSC101*, $Cm^R$ | This study |
| pXG10-*vc1461* (*cep*)* | pAL84 | 5'UTR + 20 aa of *vc1461** | Translational GFP reporter | pSC101*, $Cm^R$ | This study |
| pXG10-*vc2087* (*sucA*) | pAL72 | 5'UTR + 20 aa of *vc2087* (*sucA)* | Translational GFP reporter | pSC101*, $Cm^R$ | This study |
| pXG10-*vca0015* | pAL76 | 5'UTR + 20 aa of *vc0015* | Translational GFP reporter | pSC101*, $Cm^R$ | This study |
| pXG10-*vc0856* | pAL75 | 5'UTR + 20 aa of *vc0856* | Translational GFP reporter | pSC101*, $Cm^R$ | This study |
| pXG10-*vc2091* (*sdhC*) | pSM003 | 5'UTR + 20 aa of *vc2091* (*sdhC*) | Translational GFP reporter | pSC101*, $Cm^R$ | This study |
| pXG10-*vc2085* (*sucC*) | pSM002 | 5'UTR + 20 aa of *vc2085* | Translational GFP reporter | pSC101*, $Cm^R$ | (11) |
| pXG10-*vc2514* | pAL73 | 5'UTR + 20 aa of *vc2514* | Translational GFP reporter | pSC101*, $Cm^R$ | This study |
| pXG10-*vca0223* (*prtV*) | pNP83 | 5'UTR + 15 aa of *prtV* | Translational GFP reporter | pSC101*, $Cm^R$ | This study |
| pCMW-1 | pCMW-1 | | Control plasmid | p15A, $Kan^R$ | (12) |
| p-*cisR* | pAL41 | *cisR* | *cisR* expression plasmid | p15A, $Kan^R$ | This study |
| p-*cisR* C8G G21C G22C* | pAL85 | *cisR* C8G G21C G22C* | *cisR* C8G G21C G22C* expression plasmid | p15A, $Kan^R$ | This study |
| pEVS143 | pEVS143 | P*tac* promotor | Constitutive expression plasmid (template) | p15A, $Kan^R$ | (13) |
| pEVS-protein | pMD80 | P*tac* promotor, 5'UTR, MCS, and T1 terminator | Protein expression plasmid | p15A, $Kan^R$ | (8) |
| p-*vc0583 (hapR*) | pDD11 | *vc0583 (hapR*) | *vc0583 (hapR*) expression plasmid | p15A, $Kan^R$ | This study |
| p-*vc2614 (crp)* | pAL79 | *vc2614 (crp)* | *vc2614 (crp)* expression plasmid | p15A, $Kan^R$ | This study |
| pBAD1K-ctr | pMD004 | | Control plasmid | p15A, $Kan^R$ | (2) |
| pBAD1K-*prtV-cisR* | pAL50 | *prtV-cisR* | Inducible *prtV-cisR* expression plasmid | p15A, $Kan^R$ | This study |
| p-P*vca0224-prtV-cisR* ($P_{long}$) | pAL59 | P*vca0224-prtV-cisR* | P*vca0224-prtV-cisR* expression plasmid | p15A, $Kan^R$ | This study |
| p-*prtV-cisR* ($P_{short}$) | pAL60 | *prtV-cisR* | *prtV-cisR* expression plasmid | p15A, $Kan^R$ | This study |
| p-P*vca0224::GFP* | pAL82 | P*vca0224::GFP* | Transcriptional reporter for CisR | p15A, $Kan^R$ | This study |
| pKAS32 | pKAS32 | | suicide plasmid for allelic exchange | R6K, $Amp^R$ | (14) |

| | | | | | |
|---|---|---|---|---|---|
| pKAS32-Δ*cisR* | pAL48 | up-/downstream flanks of *cisR* | suicide plasmid for *cisR* knock-out | R6K, $Amp^R$ | This study |
| pKAS32-*vc1461(cep)::3XFLAG* | pAL55 | *vc1461(cep)::3XFLAG* | *vc1461(cep)::3XFLAG* allelic replacement | R6K, $Amp^R$ | This study |
| pKAS32-*vc2614(crp)::3XFLAG* | pKV126 | *vc2614(crp)::3XFLAG* | *vc2614(crp)::3XFLAG* allelic replacement | R6K, $Amp^R$ | This study |
| pBAD-1C-native-5'UTR-rstC | pAL89 | *rstC* | Inducible *rstC* expression plasmid | P15A, $Cm^R$ | This study |
| pKAS32- *vc0583 (hapR)::3XFLAG* | pASp017 | *vc0583 (hapR)::3XFLAG* | *vc0583 (hapR)::3XFLAG* allelic replacement | R6K, $Amp^R$ | (9) |
| pKAS32-Δ*vc2614(crp)* | pRH023 | up-/downstream flanks of *crp* | suicide plasmid for *crp* knock-out | R6K, $Amp^R$ | (8) |

### Table S3: Oligonucleotides used in this study

| Name | Sequence 5' to 3' | Description |
|---|---|---|
| **Oligonucleotides for plasmid construction** | | |
| KPO-00092 | CCACACATTATACGAGCCGA | Plasmid construction (pEVS143) |
| KPO-00196 | GGAGAAACAGTAGAGAGTTGCG | Plasmid construction (pBAD) |
| KPO-00267 | TAATAGGCCTAGGATGCATATG | Plasmid construction (pKAS32) |
| KPO-00268 | CGTTAACAACCGGTACCTCTA | Plasmid construction (pKAS32) |
| KPO-01397 | GATCCGGTGATTGATTGAGC | Plasmid construction (pEVS143, pBAD) |
| KPO-01423 | TCTAGATTAAATCAGAACGCAGAAG | Plasmid construction (pBAD) |
| KPO-01702 | ATGCATGTGCTCAGTATCTCTATC | Plasmid construction (pXG10) |
| KPO-01703 | GCTAGCGGATCCGCTGG | Plasmid construction (pXG10) |
| KPO-01952 | AGGCCTAGTTAAGGAGATATACA | Plasmid construction (pCMW) |
| KPO-01953 | GTCGACAATGAAGGGTCTTTTA | Plasmid construction (pCMW) |
| KPO-02757 | TGAGGATCCGGTGATTGATTGAGCA | Plasmid construction (pCMW) |
| KPO-02758 | AATGAAGGGTCTTTTATGATCTTAT | Plasmid construction (pDD11) |
| KPO-02759 | AAAAGACCCTTCATTCTAGTTCTTATAGATACACA | Plasmid construction (pDD11) |
| KPO-02760 | ATCACCGGATCCTCAATCCTCGCTTTGTTAT | Plasmid construction (pDD11) |
| KPO-03362 | GTTTTTATGCATAAATACTTTACATATGGATATGTACTATG | Plasmid construction (pNP83) |
| KPO-03363 | GTTTTTGCTAGCGCCTAAATCAATGGGTGTTTGAG | Plasmid construction (pNP83) |
| KPO-04168 | GACTACAAAGACCATGACGG | Plasmid construction |

| | | |
|---|---|---|
| | | (pAL55) |
| KPO-05249 | GAGATACTGAGCACATGCATGTCGCCGATTTGGCGGTTGAAATG | Plasmid construction (pSM003) |
| KPO-05250 | CCAGCGGATCCGCTAGCAATGGTCTGCAAATCTAAATTAAC | Plasmid construction (pSM003) |
| KPO-05416 | TTACTATTTATCGTCATCTTTGTAGTCG | Plasmid construction (pAL55) |
| KPO-06040 | TAGAGGTACCGGTTGTTAACGGGATGAGAGTTTTGTGGTGATC | Plasmid construction (pKV126) |
| KPO-06041 | GCGAGTGCCGTAAACCACG | Plasmid construction (pKV126) |
| KPO-06042 | CGTGGTTTACGGCACTCGCGACTACAAAGACCATGACGGT G | Plasmid construction (pKV126) |
| KPO-06043 | GACGGGTTATCGGGGCACCTATTTATCGTCATCTTTGTAGTCG | Plasmid construction (pKV126) |
| KPO-06044 | GTGCCCCGATAACCCGTC | Plasmid construction (pKV126) |
| KPO-06045 | CATATGCATCCTAGGCCTATTA CAACGCTGCTTCCTGCTACC | Plasmid construction (pKV126) |
| KPO-08333 | TCGGCTCGTATAATGTGTGGATTCTTCCTTCTCCTTCCATG | Plasmid construction (pAL41) |
| KPO-08334 | GCTCAATCAATCACCGGATCAAAAAAGGGGCACCCGAAGGTGC | Plasmid construction (pAL41) |
| KPO-08335 | GAGATACTGAGCACATGCATCATCCTTTGGGATTGGCGC | Plasmid construction (pAL42) |
| KPO-08336 | GAGCCAGCGGATCCGCTAGCAACCCCGAGTGAAAGCGTG | Plasmid construction (pAL42) |
| KPO-08384 | CGCAACTCTCTACTGTTTCTCCATGAAAACGATCAAAAAAACGCT ATTA | Plasmid construction (pAL50) |
| KPO-08385 | CTCAATCAATCACCGGATC GGCTTTTCGCATTGGCATGA | Plasmid construction (pAL50) |
| KPO-08386 | AGAGGTACCGGTTGTTAACG TGCTTGGTATTCACTCCCTG | Plasmid construction (pAL48) |
| KPO-08387 | ATCACCATCAAAGTCAAACTGTAA GGGCACCTTCGGGTGC | Plasmid construction (pAL48) |
| KPO-08388 | TTACAGTTTGACTTTGATGGTGAT | Plasmid construction (pAL48) |

| | | |
|---|---|---|
| KPO-08389 | TATGCATCCTAGGCCTATTA TTCCACGAAGCCAATCACTATTA | Plasmid construction (pAL48) |
| KPO-08418 | TAGAGGTACCGGTTGTTAACG TGCGCGTACTCGGCCTC | Plasmid construction (pAL55) |
| KPO-08419 | CCGTCATGGTCTTTGTAGTCTTTAGCCTTACGAATTAAGCCAAT | Plasmid construction (pAL55) |
| KPO-08420 | CGACTACAAAGATGACGATAAATAGTAATAGTGCTTGAGTTGTG GCTG | Plasmid construction (pAL55) |
| KPO-08421 | CATATGCATCCTAGGCCTATTACGCCGTGTTCATGGTGTC | Plasmid construction (pAL55) |
| KPO-08767 | GGAGAACGAAGAATCCACACATTATACGAG | Plasmid construction (pAL85) |
| KPO-08924 | TAAAAGACCCTTCATTGTCGAC ACAGACCTAACTAACAATTACG AT | Plasmid construction (pAL59) |
| KPO-08925 | TAAAAGACCCTTCATTGTCGACATGAAAACGATCAAAAAAACGCT ATT | Plasmid construction (pAL60) |
| KPO-08926 | TGCTCAATCAATCACCGGATCCTCAAAAAAAGGGGCACCCGAAG | Plasmid construction (pAL59, pAL60) |
| KPO-09376 | GAGATACTGAGCACATGCATGCAGGCCTTCGGGCC | Plasmid construction (pAL72) |
| KPO-09377 | GAGCCAGCGGATCCGCTAGCTGCATTGGCGCCAGCCA | Plasmid construction (pAL72) |
| KPO-09378 | GAGATACTGAGCACATGCATGCCGCGTGAAGAAACAGC | Plasmid construction (pAL73) |
| KPO-09379 | GAGCCAGCGGATCCGCTAGCTGAAATGGTCACTTCGCCTTG | Plasmid construction (pAL73) |
| KPO-09382 | GAGATACTGAGCACATGCATGAACTTTGAAGCAGCGGGC | Plasmid construction (pAL75) |
| KPO-09383 | GAGCCAGCGGATCCGCTAGCATCACGCTCGGAGGCGT | Plasmid construction (pAL75) |
| KPO-09384 | GAGATACTGAGCACATGCATCACGCTTTGCCCGCGC | Plasmid construction (pAL76) |
| KPO-09385 | GAGCCAGCGGATCCGCTAGCTGCACGCGCCAACTGATC | Plasmid construction (pAL76) |
| KPO-09649 | GCTAACAGGAGGAATTAACCATGGTTCTAGGTAAACCTCAAAC | Plasmid construction (pAL79) |
| KPO-09650 | TCGTTTTATTTGATGCCTCTAGATTATTAGCGAGTGCCGTAAAC CA | Plasmid construction |

| | | |
|---|---|---|
| | | (pAL79) |
| KPO-09800 | AAAGACCCTTCATTGTCGACTATTGATAGCATGTTTAATTTCACTG | Plasmid construction (pAL82) |
| KPO-09801 | ATATCTCCTTAACTAGGCCTGTAACAATTTATGGCTGGCGA | Plasmid construction (pAL82) |
| KPO-10106 | GGAGAACCTACATGTTTAGCTC | Plasmid construction (pAL84) |
| KPO-10107 | CATGTAGGTTCTCCTTTACCGT | Plasmid construction (pAL84) |
| KPO-10158 | TCTTCGTTCTCCTTCCATCCATTTGGG | Plasmid construction (pAL85) |
| KPO-10633 | CGCAACTCTCTACTGTTTCTCCTAGAGCTCATGCCATATTGAATT | |
| KPO-10634 | CTTCTGCGTTCTGATTTAATCTAGATTACAGTGATGGCTCAGTCAAT | |
| pBAD-ATGrev | GGTTAATTCCTCCTGTTAGC | Plasmid construction (pEVS protein) |
| pZE-Stop-XbaI | TAATCTAGAGGCATCAAATAAAACGA | Plasmid construction (pEVS protein) |
| **Oligonucleotides for Northern blot probing** | | |
| KPO-00243 | TTCGTTTCACTTCTGAGTTCGG | 5S oligoprobe |
| KPO-02077 | CCGCGAAAAGTAGGTTGTTTC | Vcr229 oligoprobe |
| KPO-02235 | AAATCCATGGAAGGAGAAGGAAG | CisR oligoprobe |
| KPO-02683 | GTGCTTAATCGTCAGCTTGTAAC | TarB oligoprobe |
| KPO-05415 | CAACGGGAGAGAAAACGGTT | Vssrna24 oligoprobe |
| KPO-08374 | ATTCTTCCTTCTCCTTCCATGG | CisR riboprobe |
| KPO-08375 | GTTTTTTTAATACGACTCACTATAGGGAGGAAAAAGGGGCACC CGAAGG | CisR riboprobe |
| **Oligonucleotides for qPCR** | | |
| KPO-00496 | GTTTGTACTTTACCGAACGC | P*vqmR* qPCR |
| KPO-02378 | GGTAACCCAGAAACTACCACTG | *recA* qPCR |
| KPO-02379 | CACCACTTCTTCGCCTTCTT | *recA* qPCR |
| KPO-07712 | CGCCAATTATGTCGGTTTC | P*vqmR* qPCR |
| KPO-08732 | GCATTTGCTAACCAAGCAC | *cep* qPCR |

| | | |
|---|---|---|
| KPO-08733 | AAGCGCAATCACCGTATC | *cep* qPCR |
| KPO-09026 | TGTCTTATTAGTCATTTATTCGATTGAT | P*cisR* qPCR |
| KPO-09027 | GGCTGGCGATAATTCATAGCAA | P*cisR* qPCR |
| KPO-09561 | GCAATCTCTACAAGACCCATT | *hfq* qPCR |
| KPO-09562 | CGATCTGACCTTGCAGTTT | *hfq* qPCR |
| **Oligonucleotides for qRT-PCR** | | |
| KPO-00584 | GCAAGGTCAGATCGAATCATTTG | *hfq* qRT-PCR |
| KPO-00585 | GGTGGCTAACTGGACGAG | *hfq* qRT-PCR |
| KPO-02745 | CATTCTTGGTGATCTCATGATAAGG | *toxT* qRT-PCR |
| KPO-02746 | CATTTACCACTTCAGAAAGGACAG | *toxT* qRT-PCR |
| KPO-02747 | AAGCAGTCAGGTGGTCTTATG | *ctxA* qRT-PCR |
| KPO-02748 | ACAAATCCCGTCTGAGTTCC | *ctxA* qRT-PCR |
| KPO-08732 | GCATTTGCTAACCAAGCAC | *cep* qRT-PCR |
| KPO-08733 | AAGCGCAATCACCGTATC | *cep* qRT-PCR |
